# ROKET: Associating Somatic Mutation with Clinical Outcomes through Kernel Regression and Optimal Transport

**DOI:** 10.1101/2021.12.23.474064

**Authors:** Paul Little, Li Hsu, Wei Sun

**Affiliations:** Biostatistics Program, Public Health Sciences Division, Fred Hutchinson Cancer Research Center, Seattle, Washington, U.S.A; Department of Biostatistics, University of Washington, Seattle, Washington, U.S.A; Department of Biostatistics, University of North Carolina at Chapel Hill, Chapel Hill, North Carolina, U.S.A

**Keywords:** Cancer, Immune response, Kernel regression, Optimal transport, Somatic point mutations, Survival

## Abstract

Somatic mutations in cancer patients are inherently sparse and potentially high dimensional. Cancer patients may share the same set of deregulated biological processes perturbed by different sets of somatically mutated genes. Therefore, when assessing the associations between somatic mutations and clinical outcomes, gene-by-gene analyses is often under-powered because it does not capture the complex disease mechanisms shared across cancer patients. Rather than testing genes one by one, an intuitive approach is to aggregate somatic mutation data of multiple genes to assess the joint association. The challenge is how to aggregate such information. Building on the optimal transport method, we propose a principled approach to estimate the similarity of somatic mutation profiles of multiple genes between tumor samples, while accounting for gene-gene similarity defined by gene annotations or empirical mutational patterns. Using such similarities, we can assess the associations between somatic mutations and clinical outcomes by kernel regression. We have applied our method to analyze somatic mutation data of 17 cancer types and identified at least three cancer types harboring associations between somatic mutations and overall survival, progression-free interval or cytolytic activity.

## Introduction

Cancer arises through somatic DNA mutations and cancer treatment often involves targeting the protein products of mutated genes. Therefore it is expected that somatic mutations may be associated with clinical outcomes. However, assessing the associations between clinical outcomes and the mutation status of each gene has met with limited success [24, 10, 12]. A likely reason is that cancer is a pathway-level disease and cancer patients may share perturbed biological processes such as a signaling pathways, but not individual genes [17]. Therefore gene-level mutations are often sparse and not shared across cancer patients. A potential solution to improve the power of somatic mutation association analysis is to aggregate the somatic mutation status of individual genes to capture the perturbation at a higher level, such as pathways or biological processes. We propose a principled approach to achieve this goal. Using the optimal transport method, we incorporate pathway or biological process annotations as well as mutual exclusive mutation patterns to define the similarity of two mutation profiles where a mutation profile is defined as a vector of mutation status for multiple genes. Then we use such similarity metrics to perform kernel regression against clinical outcomes.

A naive way to define the distance/similarity of two mutation profiles is to apply a standard distance metric (e.g., Euclidean distance) without considering additional information such as gene annotations. Such a generic distance definition implicitly assumes each gene in the mutation profile independently contributes to the distance and does not exploit dependence across genes. Our task is thus two-fold: first, quantify the dependencies or similarities across genes; second, transfer such 75 similarity across genes into the similarity across mutation profiles of tumor samples in a principled way.

We will explore three methods to quantify the similarity of two genes: whether they share similar Gene Ontology annotations [20], whether they belong to the same set of pathways, and whether their mutation patterns are mutually exclusive, which implies the mutation of these two genes play redundant functional roles in tumor growth.

Next we transform such gene-gene similarity to sample-sample distance using optimal transport. Optimal transport involves the idea of “transporting mass” or mapping from one space or distribution to another by minimizing the total cost given the known costs associated with all possible choices of transportation. The optimal transport framework has been applied to a wide range of fields and applications such as statistics, natural language processing, business/economics, image analysis, and domain mapping [22, 13]. For our application, we seek to transport the somatic mutational profiles between two tumor samples and use the cost of optimal transport to measure their distance. The cost of each transportation plan can be calculated using the above mentioned gene-gene similarity. For instance, consider two samples (sample 1 and 2) and two genes (denoted by A and B). Suppose sample 1 has mutation in gene A and sample 2 has mutation in gene B. If the two genes are not related, the cost to transport mutation from gene A to gene B is high. In contrast, if the two genes belong to the same pathway, the cost of transport is low. Therefore the cost of transport can be used as a distance measure between samples informed by the gene-gene relationship. The resulting sample-sample distance matrix can then be transformed into a kernel for kernel regression.

There are several challenges or practical issues to implement optimal transport (OT) method for somatic mutation data. First, gene annotation may not be accurate or complete, and thus it is desirable to aggregate multiple types of annotations to define gene-gene similarity. Second, OT has associated penalty parameters to regulate how the transported mass are scaled and distributed. In somatic mutation data, the tumor mutation burden (TMB), i.e., the number of mutated genes per sample, often varies across samples. Quantifying the cost of transportation between a sample with many mutations and another with few mutations requires exploration of a range of penalty parameter values. Third, a practical issue is how to select a subset of genes for OT. Our work addresses all these challenges and issues by using an omnibus test strategy [9] to aggregate the kernels calculated using different set of parameters.

We have developed an R package ROKET (**R**egression **O**perated by **KE**rnels and **T**ransport) to implement OT of somatic mutations and to perform kernel association testing. We performed simulations to demonstrate the power gains our OT-based approaches in association analyses over gene-by-gene association or kernel regression using a naive distance definition by the Euclidean distance. Then we applied our method to conduct a pan-cancer analysis of 17 cancer types and revealed somatic mutations are significantly associated with different outcomes in three cancer types after accounting for the effects of other covariates such as tumor stage and mutation burden.

## Methods

### Notation and model

Consider *N* tumor samples. For each sample *i* = 1*,…, N*, we have somatic mutation status on *G* genes, denoted by a *G* × 1 vector ***Z**_i_* = (*Z_i_*_1_*,…, Z_iG_*)^T^, where *Z_ig_* = 1 if the *g*th gene has at least one non-synonymous mutation and 0 otherwise, and T denote the transpose operator. We also have *p* baseline covariates denoted by a *p* × 1 vector ***X**_i_*. For survival outcomes, let 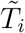 and *C_i_* denote event and censoring time, respectively, such that we observe 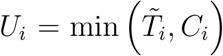 and 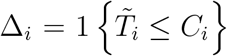. We assume *C_i_* is independent of 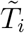 conditioning on ***X**_i_* and ***Z**_i_*. For non-survival outcome, we denote it by *Y_i_* with expectation *E* [*Y_i_|**X**_i_, **Z**_i_*].

We model survival outcomes using the semi-parametric kernel machine Cox proportional hazards model [4]. The hazard function is characterized as

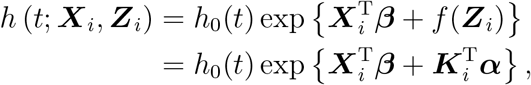

where *h*_0_(*t*) is an unspecified baseline hazard function. We adjust for baseline covariates through a parametric component 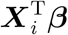 and the high-dimensional covariate through the non-parametric component *f* (***Z**_i_*). We assume *f* (·) is generated by a positive semi-definite function *K*(·, ·) such that f(·) lies in the reproducing kernel Hilbert space 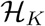. With the representer theorem [7], we have 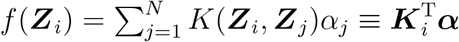 for parameters **α** and kernel matrix ***K*** = *K*_(*i,j*)_ = [*K*(***Z**_i_, **Z**_j_*)]. Through the mixed effects model framework, we consider *f* (***Z**_i_*) as a random effect where vector (*f* (***Z***_1_)*,…, f* (***Z**_N_*)) has mean 0 and variance *τ**K***. Kernel significance testing involves *H*_0_ : *f* (***Z***) = ***Kα*** = 0 vs. *H_A_* : *f* (***Z***) ≠ 0 which is equivalent to testing *H*_0_ : *τ* = 0 vs. *H_A_* : *τ* ≠ 0 [11]. The score test is used with the benefit of avoiding estimation of the full mixed effects model, which can be computationally intensive especially for non-linear outcome. The asymptotic p-value can be calculated by inverting the characteristic function. In the situation of small sample size or testing of multiple kernels, a permutation-based p-value may be preferred to better approximate the null distribution. This can be obtained by permuting the null model’s residuals.

For binary and continuous outcomes, we have

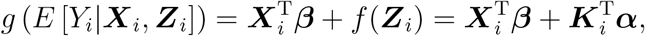

where *g*(·) is a known link function, e.g., linear link for continuous outcome and logit link for binary outcome.

The following sections discuss how ***K*** is obtained from an *N* × *N* distance matrix ***D*** = [*D_ij_*] where *D_ij_* ≡ *D*(***Z**_i_, **Z**_j_*). The matrix ***D*** is calculated by optimal transport of gene mutation statuses given pre-computed gene-gene similarity.

### Optimal Transport (OT)

The goal of OT is determining how to optimally transport mass. In our case, we want to map the gene mutation profile from one sample to another sample. To formulate OT, let ***P*** = [*P_gh_*] denote a transport plan, which is a *G* × *G* matrix where *P_gh_* represents how much mass is transported between elements *g* and *h*. Let ***W*** = [*W_gh_*] denote a known *G × G* cost matrix with *W_gh_* representing the cost to transport mass between elements *g* and *h* and *W_gg_* = 0 as there is no cost to transport mass between identical elements. *W_gh_* is defined as one minus the similarity between the *g*th and *h*th genes. The total cost to transport mass between two samples ***Z**_i_* and ***Z**_j_* is defined as

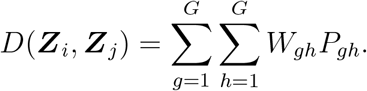

With this framework, the goal is to find ***P*** such that *D*(***Z**_i_, **Z**_j_*) is minimized. The resulting minimization can then be used to characterize the “distance” between ***Z**_i_* and ***Z**_j_*, which can be further used for kernel regression or unsupervised clustering. Note that *D*(***Z**_i_, **Z**_i_*) = 0 since *W_gg_* ≡ 0. There are two types of optimal transport problems: balanced OT where the mass of two input vectors are the same or pre-normalized, and unbalanced OT otherwise.

To solve for ***P*** given ***W*** and ***Z***_1_*,…, **Z**_N_*, we must consider several questions. First, how to balance the computational efficiency and the accuracy of the results by adjusting the tuning parameters. Second, the mutation burden varies across tumor samples. Should the optional transport account for such variation of mutation burden or is it better to pre-normalize the mutation data so that all the samples have the same mutation burden? A related question is that the somatic mutation data are often sparse. For example, a sample may have mutation in only 1% of the ~ 20,000 genes. Could we exploit such sparsity to improve computational efficiency?

To improve the computational efficiency, a well-studied and popular solution is to regularize the objective function by adding a term *ϵ* Σ*_g,h_ P_gh_* log (*P_gh_*), where *ϵ* is a penalty parameter. This entropic regularization follows the maximum-entropy principle, and the regularization transforms a linear programming problem into a strictly convex problem that can be solved quickly with multi-threaded code [5]. A smaller *ϵ* leads to more accurate solutions and a larger *ϵ* improves computational efficiency. An appropriate choice of *ϵ* depends on the problem of interest. When the problem of interest is to make predictions, *ϵ* can be chosen by cross-validation. However, our interest is to assess associations. We choose the largest *ϵ* while it still guarantees the distance between one sample and itself is close to 0. More specifically, we determined *ϵ* such that max*_i_ D_ii_* is < 0.001.

To consider different options to handle the variation of mutation burden, we need to first introduce two types of OT: balanced OT and unbalanced OT. Balanced OT refers to pre-normalizing each sample’s mass to a constant and then constraining ***P*** such that *Z_ig_* = Σ*_g_ P_gh_* and *Z_jh_* = Σ*_h_ P_gh_* by applying Lagrange multipliers to the objective function. With *ϵ* = 0, the “UV” or MODI method [6] finds an exact solution for ***P***. With *ϵ* > 0, taking partial derivatives of the objective function leads to alternating updating equations

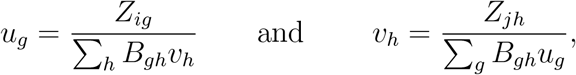

where 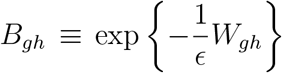 and *P_gh_* = *u_g_B_gh_v_h_*. The vectors *u* = (*u_1_,…, u_G_*) and *v* = (*v*_1_*,…, v_G_*) represent how much one input vector’s elements have to be transported to and arrive at the other vector. Notice that if any *Z_ig_* = 0 (or *Z_jh_* = 0), the corresponding *u_g_* = 0 (or *v_h_* = 0), indicating that *P_gh_* = 0. This means when performing the OT, we do not need to consider the genes that are not mutated. For example, sample 1 has mutations in 100 out of 20,000 genes and sample 2 has mutations in 200 out of 20,000 genes. We just need to focus on the 100 × 200 transportation matrix instead of the 20, 000 × 20, 000 transportation matrix. This strategy utilizes the sparsity of the somatic mutation data to reduce computational costs.

Unbalanced OT, with *ϵ* > 0, avoids the step of pre-normalizing sample masses. So the Lagrange constraints within the objective function need to be generalized. A popular solution is to replace the Lagrange terms with generalized Kullback-Leibler divergence defined as

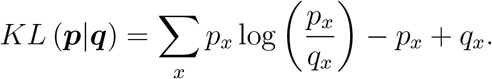

For shorthand notation, let 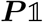 and 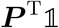 denote the row and column sums of ***P***, respectively.

The unbalanced OT objective function with regularization is defined as

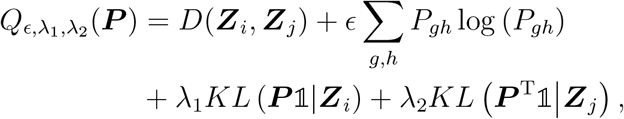

where *ϵ*, *λ*_1_, and *λ*_2_ are all tuning parameters. The values of *λ*_1_ and *λ*_2_ control how much ***P***’s marginalized mass mimics the mass of ***Z**_i_* and ***Z**_j_*, respectively. To solve the above unbalanced OT objective function, we arrive at the alternating updating equations

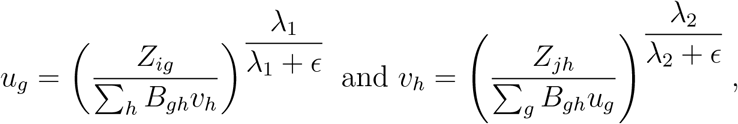

where *B_gh_* and *P_gh_* follow the same definitions as above. In our exploratory analysis, setting *λ*_1_ ≠ *λ*_2_ does not lead to apparently better performance but dramatically increases computational time. Therefore, we restricted simulations and analyses to *λ*_1_ = *λ*_2_ = *λ* and performed the association analysis using balanced OT as well as unbalanced OT with *λ* ∈ {0.5, 1.0, 5.0}, and then combined the results across kernels using an omnibus test approach [9], i.e., choosing the kernel that leads to the strongest association and assessing its significance while accounting for the search over multiple kernels.

### Gene Similarity

OT requires a known or pre-calculated cost matrix ***W***. In our work, this cost to transport mass between genes A and B is one minus the similarity between genes A and B. The gene similarities were derived from three sources: gene ontology (GO), canonical pathways, and mutually exclusive (ME) analysis, which we refer to as GO-based, Path-based, and ME-based, respectively. The GO-based approach characterizes the similarity of two genes by the similarities of their associated GO Terms. Since GO terms are organized by a directed acyclic graph, the similarity of GO terms can be calculated using their locations in the graph [21, 20]. We evaluated GO-based similarity using R v4.0.2 with packages GOSemSim v2.14.2 and obtained GO annotations (biological process) from R packages org.Hs.eg.db v3.11.4 and AnnotationDbi v1.50.1. For two genes annotated with *g*_1_ and *g*_2_ GO terms, respectively, their GO terms’ similarity forms a matrix of size *g*_1_ × *g*_2_. Following the approach of Wang et al. [20], we reduced such a distance matrix to a scalar by “best-match average”, which calculates the average of all maximum similarities on each row and column of this similarity matrix.

Unlike the GO-based approach with an underlying tree structure, there is no natural definition of similarity between pathways. We propose an approach to define gene-gene similarity using pathway information (PATH approach), by quantifying whether two genes likely share their pathway annotation by chance. We used the reactome pathway annotation obtained from Molecular Signatures Database version 7.2. Starting with the reactome file containing pathways and their associated genes, we characterized gene similarities as follows. Let *G* be the total number of annotated genes. First, given a pair of genes, we identified the set of shared pathways. Next, we determined *S*, the number of genes sharing the same set of pathways as the original gene pair. Then, we calculated the probability of two randomly selected genes sharing the same set of shared pathways as the original gene pair by a hyper-geometric distribution. A small probability indicates the pathway shared between the two genes is unlikely to happen by chance. Finally, we subtracted this probability from one to arrive at a gene similarity.

The final candidate for gene similarity rests on the idea of mutually exclusive mutations. If mutation in one of two genes is enough to perturb a pathway that facilitates tumor growth, it would be rare to observe mutations in both genes in a tumor genome, due to the randomness of somatic mutation accumulation [3]. Mutually exclusive (ME) genes may be tumor-specific and thus gene-gene relationships are allowed to vary across cancer types. For the ME-based approach, we initially calculated one-sided Fisher exact test p-values for each pair of genes to score the association between gene mutations but soon realized that a handful of frequently mutated genes confounded the notion of mutually exclusive genes. Instead, we ran logistic regression with the first gene’s mutation status as outcome and the second gene’s mutation status as predictor, while adjusting for log(TMB) and the log(TMB) squared, and then tested the association using the one-sided test. We repeated this process with the second gene as the outcome and took the geometric mean of these two p-values to construct a symmetric matrix to score all gene pairs. Smaller scores mean smaller p-values, hence stronger patterns of mutual exclusivity, and thus larger gene-gene similarity. The distribution of these scores was concentrated between 0.5 and 1.0. Only a handful of the scores were sufficiently small, e.g., < 0.0001. So we transformed the scores by negative log(score), and normalized by the 99th percentile rather than the maximum for robustness. Any newly transformed values greater than one were set to one. This procedure transformed the original scores into gene-gene similarities in the range of 0 to 1.

### Kernel calculation

Many methods have been proposed to perform kernel regression association analysis, including KSPM [18], MiRKAT/MiRKAT-S [23, 15], and OMiSA [8]. KSPM currently only implements linear models and its implementation involves estimating penalized kernel parameters under the full model which is not necessary for hypothesis testing. We chose to proceed with the MiRKAT method and R package v1.1.4 which performs kernel regression with ordinary least squares (OLS), logistic regression, and Cox proportional hazard models with MiRKAT-S.

We calculate *K*(***Z**_i_, **Z**_j_*) as a function of pairwise distance *D_ij_* = *D*(***Z**_i_, **Z**_j_*). For our analysis, *W_gh_* ranges between zero and one and denotes the cost (dissimilarity) of transporting mass between genes *g* and *h* where a smaller value of *W_gh_* indicates the pair of genes are relatively similar, e.g., they belong to the same set of pathways.

Finally, to transform a distance matrix ***D*** = [*D_ij_*] into a positive semi-definite kernel matrix ***K***, we use

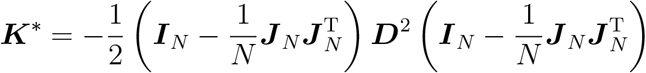

where ***D***^2^ is the element-wise squared distance matrix; ***I**_N_* and ***J**_N_* denote a *N × N* identity matrix and a vector of *N* ones, respectively. This transformation is known as double centering or centering the rows and columns of ***D*** where the original distance can be recovered by the kernel distance: 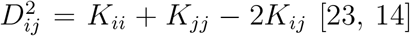 [23, 14]. The matrix ***K***^*^ is not guaranteed to be positive semi-definite and thus the final kernel matrix is calculated by performing eigenvalue decomposition of ***K***^*^ and replacing negative eigenvalues with 0’s and recalculating to obtain a positive semi-definite kernel matrix ***K***.

Our method involves several alternative choices or tuning parameters, such as the choices of gene-gene similarity or the value of parameter *λ*. Another choice in practice is how to filter out genes to be used. Some rarely mutated genes are more likely to be passenger mutations that are less likely associated with clinical outcomes. We consider a few cutoffs to select genes that are mutated in a certain number of tumor samples. Each of those choices/tuning parameters leads to a kernel. We combine the results of multiple kernels by an omnibus test [9]. Briefly, the kernel that leads to the minimum p-value is selected and the statistical significance is assessed by permuting the null model’s residuals and calculating the proportion of permutations where the permutation-based test statistic is more extreme than the observed test statistic.

### Software Implementation

At the time of completing this project and to our knowledge, there was no existing R package to perform unbalanced OT and thus we developed a stand-alone R package **R**egression **O**perated by **KE**rnels and **T**ransport (ROKET) to run balanced or unbalanced OT across a set of samples with one or multiple threads. In addition, we included the implementation of the permutation-based omnibus test using C++ in our package, which substantially improves the computational efficiency of permutation test. Our implementation of permutation test runs at least 10 times faster than existing packages.

## Simulation Study

The aims of the simulation are to verify our software implementation, and more importantly, to compare the performance of a gene-based approach (gene-by-gene associations), kernel regression using Euclidean distance, and kernel regression using OT. For OT-based approaches, we explored the performance of unbalanced vs. balanced OT. For unbalanced OT, we also explored the choice of penalty parameter *λ*.

We started by specifying the number of genes and pathways. Given *G* genes and *Q* pathways, we randomly assigned genes to pathways. Next, we simulated gene-gene similarities based on whether genes were within a pathway. Specifically, we simulated the similarity between the *g*th and *h*th genes, denoted by 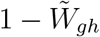, by sampling from a uniform distribution on [*δ*, 1] for 0.5 ≤ *δ* < 1 if they belonged to the same pathway, and a uniform distribution on [0, 1 − *δ*] otherwise. We added perturbation to the similarity matrix by generating 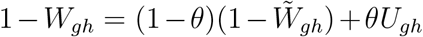 where *U_gh_* is sampled from uniform [0,1]. This represented the observed gene-gene similarity matrix. In our simulations, we specified *G* = 500, *Q* = 25, *δ* = 0.5, and *θ* = 0.2.

For baseline covariates ***X**_i_* with *i* = 1*,…, N*, we included the intercept, a binary variable generated by a Bernoulli distribution, a four level categorical variable generated by a discrete uniform distribution, and an age covariate sampled from the discrete uniform between integers 20 and 80. These variables served to represent baseline variables gender, tumor stage, and age at diagnosis. We set sample size *N* = 200.

Before simulating mutated genes, we first simulated mutated pathways per sample. Let ***R**_i_* = (*R*_*i*1_*,…, R_iQ_*) denote the *i*th sample’s mutated pathway status, where *R_iq_* = 1 if the *q*th pathway was mutated and 0 otherwise. We designed four scenarios (Table 1). In scenarios 1 and 2, the number of mutated pathways was sampled randomly from 9, 10, or 11. In scenarios 3 and 4, the number of mutated pathways was randomly sampled from integers ranging from 5 to 20. This was designed to examine how the variation of pathway burden across samples affected OT performance. For a sample with *A* pathways mutated, ***R**_i_* was generated by randomly selecting *A* elements as ones and setting the remaining *Q − A* elements as zeros. This was repeated for all *N* samples.

**Table 1:** Four simulation scenarios. PB = number of mutated pathways. TMB = number of mutated genes.

| Scenario | Varied PB | TMB ∝ PB |
| --- | --- | --- |
| 1 | NO | YES |
| 2 | NO | NO |
| 3 | YES | YES |
| 4 | YES | NO |

To simulate gene mutation statuses ***Z**_i_*, for scenarios 1 and 3, the sample TMB was three times the number of mutated pathways. For scenarios 2 and 4, the sample TMB was drawn from the discrete uniform ranging from the number of mutated pathways plus 5 to a maximum of 80. The purpose of this design was to allow TMB to be strongly or weakly correlated with the number of mutated pathways. We then sampled TMB mutated genes from the genes associated with the mutated pathways. If by chance no gene was selected from a mutated pathway, we purposely sampled one gene from that pathway.

With baseline covariates ***X**_i_* and mutated pathways ***R**_i_*, we simulated continuous and survival outcomes. The continuous outcome was generated from a linear model

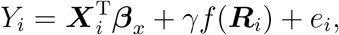

where *e_i_* is mean zero Gaussian distributed with standard deviation 0.40 and ***β**_x_* = (−5.0, 0.5, 1/3, 2/3, 1, −1)^T^. The time-to-event was generated under the Cox proportional hazards model with a Weibull distribution with shape parameter of 2 for the baseline hazard function. The time-to-event was generated by

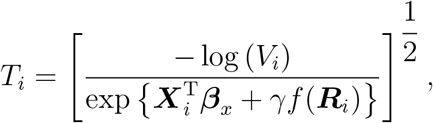

where *V_i_* was sampled from Uniform[0,1]. The scalar *γ* in both models takes values 0.0, 0.1,…, 0.9, 1.0, and *f* (***R**_i_*) is the linear summation of 1-way, 2-way, 3-way, and 4-way pathway interactions. Coefficients of the *d*-way interactions were sampled from a mean zero Gaussian with standard deviation of 0.40 multiplied by a Bernoulli with success probability 0.5^*d*−1^. Censoring times were generated independent of event time from Uniform[0, *m*], where *m* was tuned to yield censoring proportions of 75%, 50%, 25%, and 0%. High censoring proportions were included to reflect censoring proportions observed in the real data.

To evaluate the performance (power and type I error) of a kernel-based approach, we tallied whether its p-value was less than a pre-specified significance level *α*. For a gene-based approach, we tested the genes that were mutated in at least 20 samples. To ensure testing of the same null hypothesis, i.e., no overall association with the outcome, we calculated the p-value for the gene-based testing as follows. We first calculated the minimum p-value of all the tested genes. Then we calculated the permutation p-value for the minimum p-value to account for multiple testing across multiple genes. When permuting the data, we permuted the sample order of the gene-sample mutation matrix to preserve the gene-gene correlation structure and calculated the minimum p-value for the permuted data set. The permutation p-value is the proportion of permutations whose minimum p-values are smaller than the observed minimum p-value. All rejections were based on *α* = 0.05.

### Performance

We ran 200 replicates under each simulation scenario and considered both balanced and unbalanced OT. As described in Section 2.2, we set *ϵ* = 0.001. While *λ* could affect the distance calculation, to avoid an exhaustive grid search, we chose a wide range of the values from 0.5 to 5.0 to explore any trends in type I error, power, and computational run time. The omnibus OT approach was included alongside gene-based and Euclidean associations approaches. The results are shown in Figure 1. There was no evidence of inflated type I error for all the approaches.

**Figure 1:**
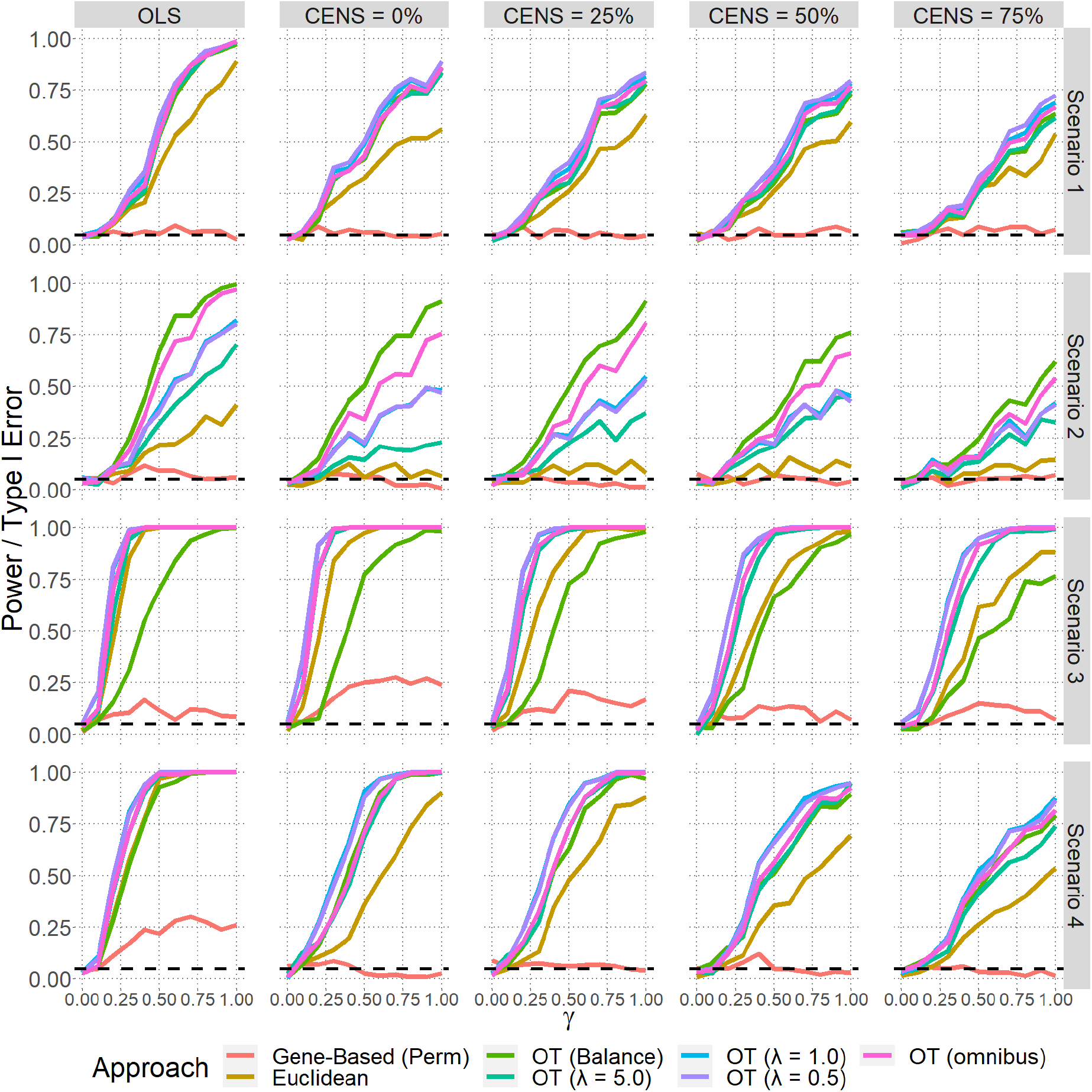
Power/type I error comparison of gene-based, Euclidean distance, balanced and unbalanced (*λ* < ∞) OT, and omnibus for association testing of somatic mutation. Y-axis corresponds to the proportion of replicates rejecting the null hypothesis. X-axis corresponds to the regression coefficient *γ* for the overall strength of the association of the mutated pathways with the outcome. Continuous, denoted by OLS, and survival outcome models are presented.

For the continuous outcome, kernel regression using either Euclidean and OT kernel were considerably more powerful than gene-based association testing (Figure 1). Under scenarios 1 and 2, where the number of mutated pathways (pathway burden, i.e., PB) was relatively uniform across samples, all OT approaches that incorporated the pathway information had greater power than the Euclidean distance-based power. The balanced OT had comparable or greater power than the unbalanced OT, despite the variation of TMB across samples. On the other hand, when the PB varied, balanced OT suffered power loss, especially in Scenario 3, because the pre-normalized input vectors did not capture the underlying PB variability. Unbalanced OTs were at least as powerful as or more powerful than the Euclidean distance, suggesting that incorporation of pathway information increases the power for detecting the associations. Regarding the three values of the tuning parameter *λ* for unbalanced OT, there was no single value that outright outperformed the rest. The omnibus OT test that combines balanced OT and three unbalanced OT distances showed greater power than the Euclidean approach. The results for survival outcome were consistent with the results for continuous outcome. Unsurprisingly, a higher censoring rate led to decrease in power.

To explore the performance of individual genes without multiple testing adjustment visualized by boxplots, type I error was maintained under the null and subtle increases in median power were observed across tested genes (Web Figure 1). Generally, the (unbalanced) OT approaches have better performance than the Euclidean approach. A key observation is that unbalanced OT appears to be robust under all scenarios whereas balanced OT is sensitive to the distribution of mutated pathways, which can not be directly observed.

## Pan-cancer Application

The Cancer Genome Atlas (TCGA) project provides researchers with multi-omic data spanning 33 cancer types collected from multiple institutions. These omic data characterize the molecular portrait of each cancer type in order to improve cancer diagnosis, treatment, and prevention [19]. For real data application, we applied our method and workflow on TCGA somatic mutation data from seventeen cancer types with sample size larger than 150 and at least 50 observed non-censoring events per survival outcome for overall survival (OS), progression-free interval (PFI) or disease-specific survival (DSS). We also considered immune cytolytic activity (CYT), which quantifies immune activity based on the geometric mean of the expression (transcripts per million) of two genes: *GZMA* and *PRF1* [16]. We examined CYT since both somatic mutation burden and individual mutations might be correlated with immune activity, which in turn may be associated with response to immunotherapy. We modeled log-transforming CYT as a continuous outcome by linear regression model.

### Workflow Overview

To arrive at the dataset for analysis per cancer type, we began with the Genomic Data Commons (GDC) resource to obtain somatic point mutation calls by MuTect2 using hg38 aligned reads. We obtained tumor purity and ploidy, demographic and clinical data from the GDC’s PanCanAtlas publications webpage (see links in the Data Availability Section). BWA version, sequencing center, and exome capture kit variables were obtained from Buckley et al. (2017) [2]. We excluded variants that failed to pass GDC filtering criteria including ‘ndp’ (normal depth filter), ‘nonexonic’, ‘exac’ (germline variant), ‘bitgt’ (targeted locus), ‘gdc pon’ (panel of normals), and ‘wga’ (whole genome amplication artifact) flags. We only included non-synonymous variants in our analysis. To obtain gene mutation information per sample, we defined a gene as mutated if it harbored at least one non-synonymous somatic point mutation.

### Pan-cancer Analyses

So many aspects were not known a priori. First, while genes mutated more frequently in a cancer population are more likely to be functionally relevant to cancer etiology, it is not clear what is an appropriate threshold to select those genes. Second, which OT penalty would best serve our problem? Third, which of the three candidates for characterizing gene similarity is the best? We evaluated all kernels corresponding to these three choices by an omnibus test. The Euclidean-based omnibus test, which considers kernels for different mutation frequency cutoffs, would serve as the benchmark against our OT approaches. If the omnibus Euclidean or OT-based kernel test was significant (p-value < 0.05) for any outcome of a cancer type, we would summarize the individual kernel test results to assess any consistent trends at various gene mutation thresholds between Euclidean and OT methods.

For each of the 17 TCGA cancer types, given a gene mutation frequency cutoff, we generated 12 OT-based distance matrices (combinations of 3 gene similarities and 4 OT approaches). We also considered various thresholds for mutation frequency such that we only included genes that mutated in at least a certain number of samples. One limit of incorporating more genes in the OT calculation was the growing number of genes missing annotation. In these situations, pairwise gene similarities were set to zero (dissimilarity set to one), treating these genes as equidistant features from each other. The number of uninformed gene pairs for GO-based and PATH-based OT distance calculations can be seen in Web Figure 2. The ME approach is not informative for genes that are mutated in a very small number of individuals. Thus when calculating ME-based similarity, we defined it for the top 300 mutated genes per cancer type and set all other gene similarities to zero. The number of genes selected based on mutation frequency varies greatly across cancer types (Table 2). For colon (COAD) and rectal (READ) cancer, denoted by COAD for short, a subset of tumors (~ 17%) were hypermutated (i.e., with a large number of mutations) and we ran separate analyses for all tumors or only non-hypermutated (nHM) tumors with TMB less than 10^2.5^, based on the log10 TMB bimodal distribution (Web Figure 3). For a handful of cases, samples might have no genes that met the mutation frequency threshold and in those cases we simply excluded them for all analyses. An example of our distance-calculation workflow starting from somatic mutations and incorporating ME-based gene annotations is presented (Figure 2). Visually, Euclidean-based distances revealed at least three clusters of samples (Figure 2(C)). In contrast, OT combined with gene-genes similarity by mutual exclusivity reveals more clusters and higher between-cluster similarities.

**Table 2:**
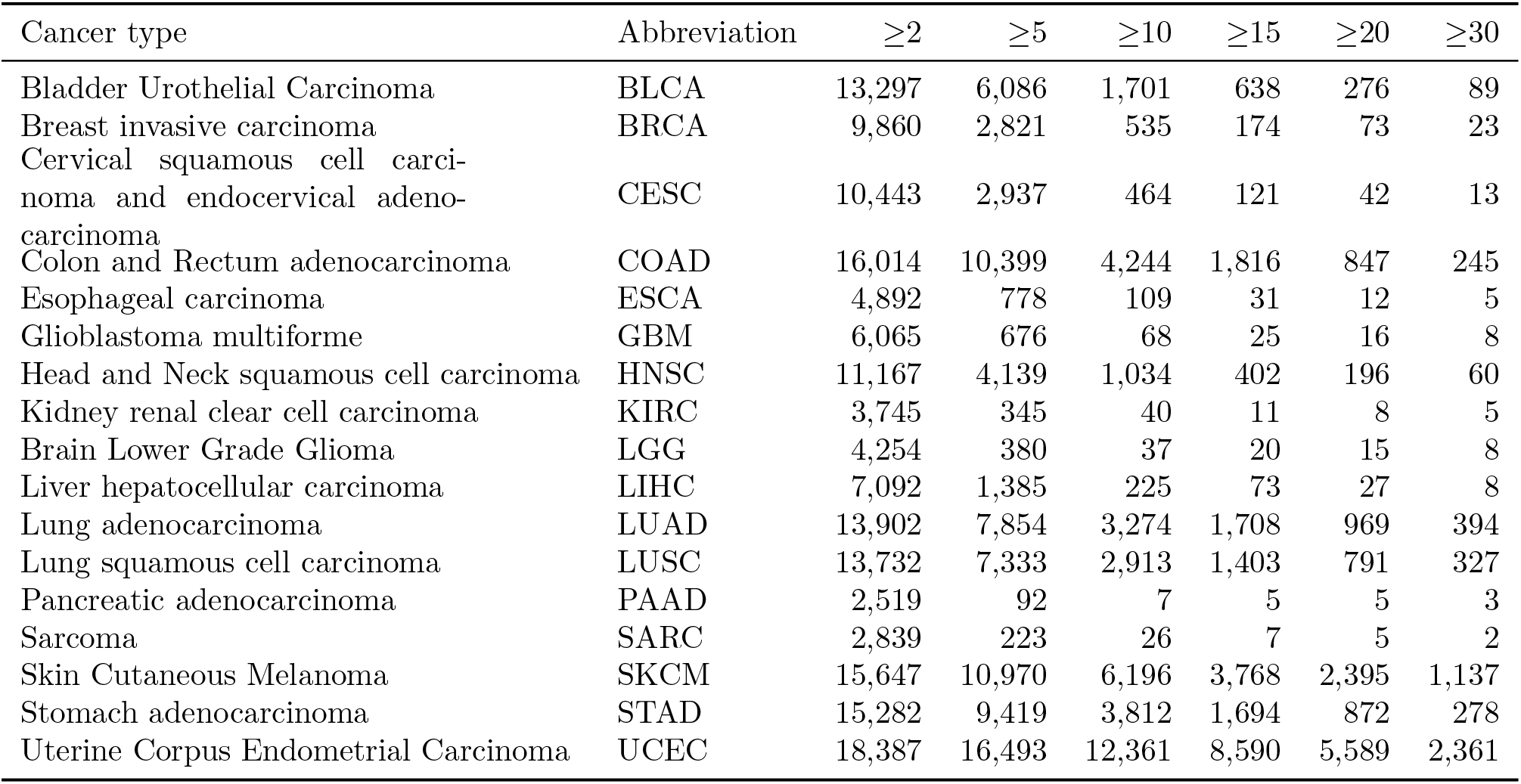
The number of genes at various mutation frequency cutoffs for each cancer type.

| Cancer type | Abbreviation | $\geq 2$ | $\geq 5$ | $\geq 10$ | $\geq 15$ | $\geq 20$ | $\geq 30$ |
| --- | --- | --- | --- | --- | --- | --- | --- |
| Bladder Urothelial Carcinoma | BLCA | 13,297 | 6,086 | 1,701 | 638 | 276 | 89 |
| Breast invasive carcinoma | BRCA | 9,860 | 2,821 | 535 | 174 | 73 | 23 |
| Cervical squamous cell carcinoma and endocervical adenocarcinoma | CESC | 10,443 | 2,937 | 464 | 121 | 42 | 13 |
| Colon and Rectum adenocarcinoma | COAD | 16,014 | 10,399 | 4,244 | 1,816 | 847 | 245 |
| Esophageal carcinoma | ESCA | 4,892 | 778 | 109 | 31 | 12 | 5 |
| Glioblastoma multiforme | GBM | 6,065 | 676 | 68 | 25 | 16 | 8 |
| Head and Neck squamous cell carcinoma | HNSC | 11,167 | 4,139 | 1,034 | 402 | 196 | 60 |
| Kidney renal clear cell carcinoma | KIRC | 3,745 | 345 | 40 | 11 | 8 | 5 |
| Brain Lower Grade Glioma | LGG | 4,254 | 380 | 37 | 20 | 15 | 8 |
| Liver hepatocellular carcinoma | LIHC | 7,092 | 1,385 | 225 | 73 | 27 | 8 |
| Lung adenocarcinoma | LUAD | 13,902 | 7,854 | 3,274 | 1,708 | 969 | 394 |
| Lung squamous cell carcinoma | LUSC | 13,732 | 7,333 | 2,913 | 1,403 | 791 | 327 |
| Pancreatic adenocarcinoma | PAAD | 2,519 | 92 | 7 | 5 | 5 | 3 |
| Sarcoma | SARC | 2,839 | 223 | 26 | 7 | 5 | 2 |
| Skin Cutaneous Melanoma | SKCM | 15,647 | 10,970 | 6,196 | 3,768 | 2,395 | 1,137 |
| Stomach adenocarcinoma | STAD | 15,282 | 9,419 | 3,812 | 1,694 | 872 | 278 |
| Uterine Corpus Endometrial Carcinoma | UCEC | 18,387 | 16,493 | 12,361 | 8,590 | 5,589 | 2,361 |

**Figure 2:**
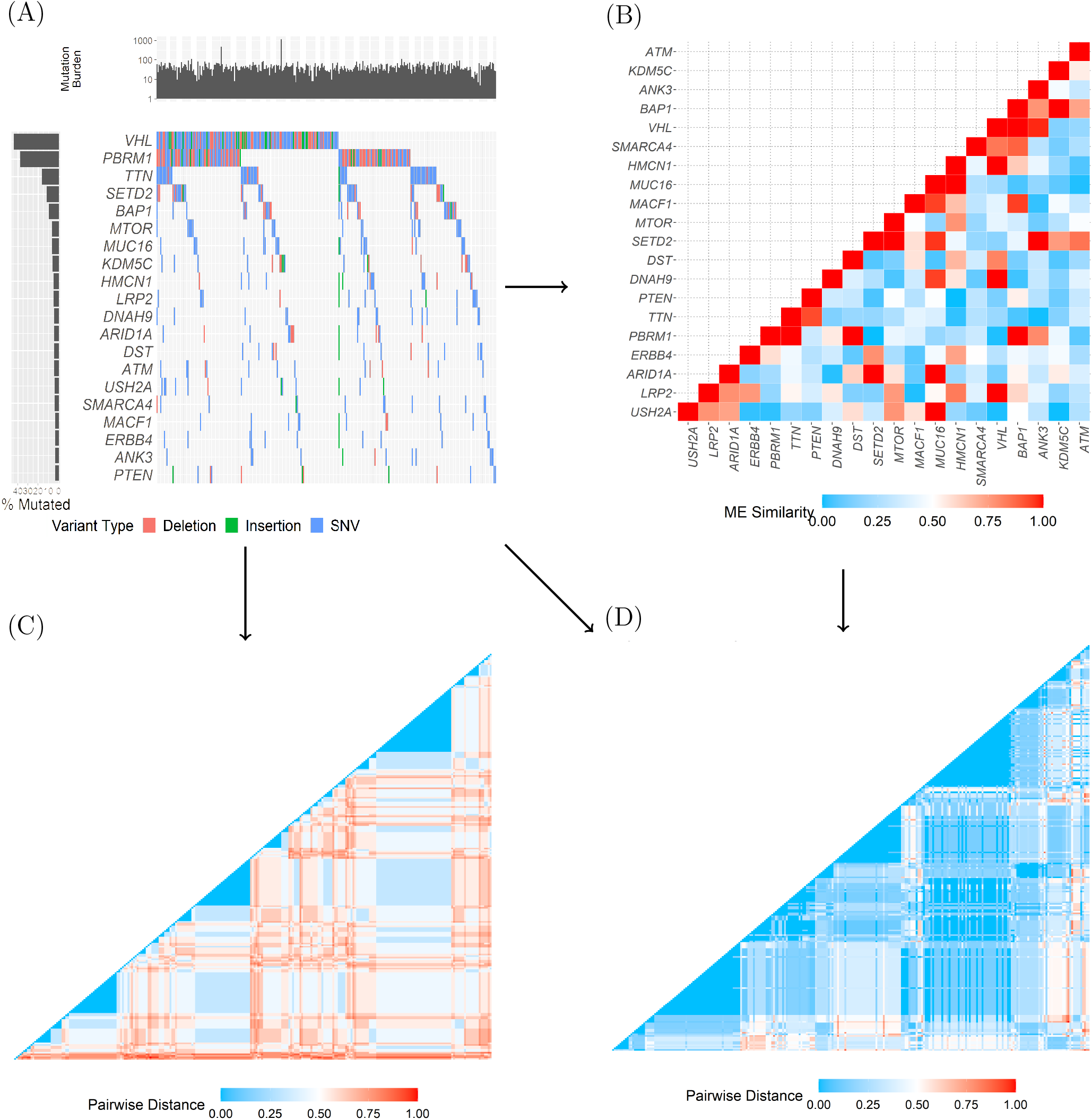
Workflow from somatic point mutations to distance matrices for TCGA’s kidney renal clear cell carcinoma. (A) A plot of samples by the top 20 mutated genes with barplots of tumor mutation burden and mutation frequency. (B) A gene-gene similarity matrix calculated based on our mutual exclusivity approach. (C) Euclidean pairwise distance heatmap. (D) Optimal transport-based pairwise distance heatmap using *λ* = 0.5 and mutual exclusivity based gene similarity. Distance matrices were independently ordered by hierarchical clustering and then scaled to range from 0 to 1.

**Figure 3:**
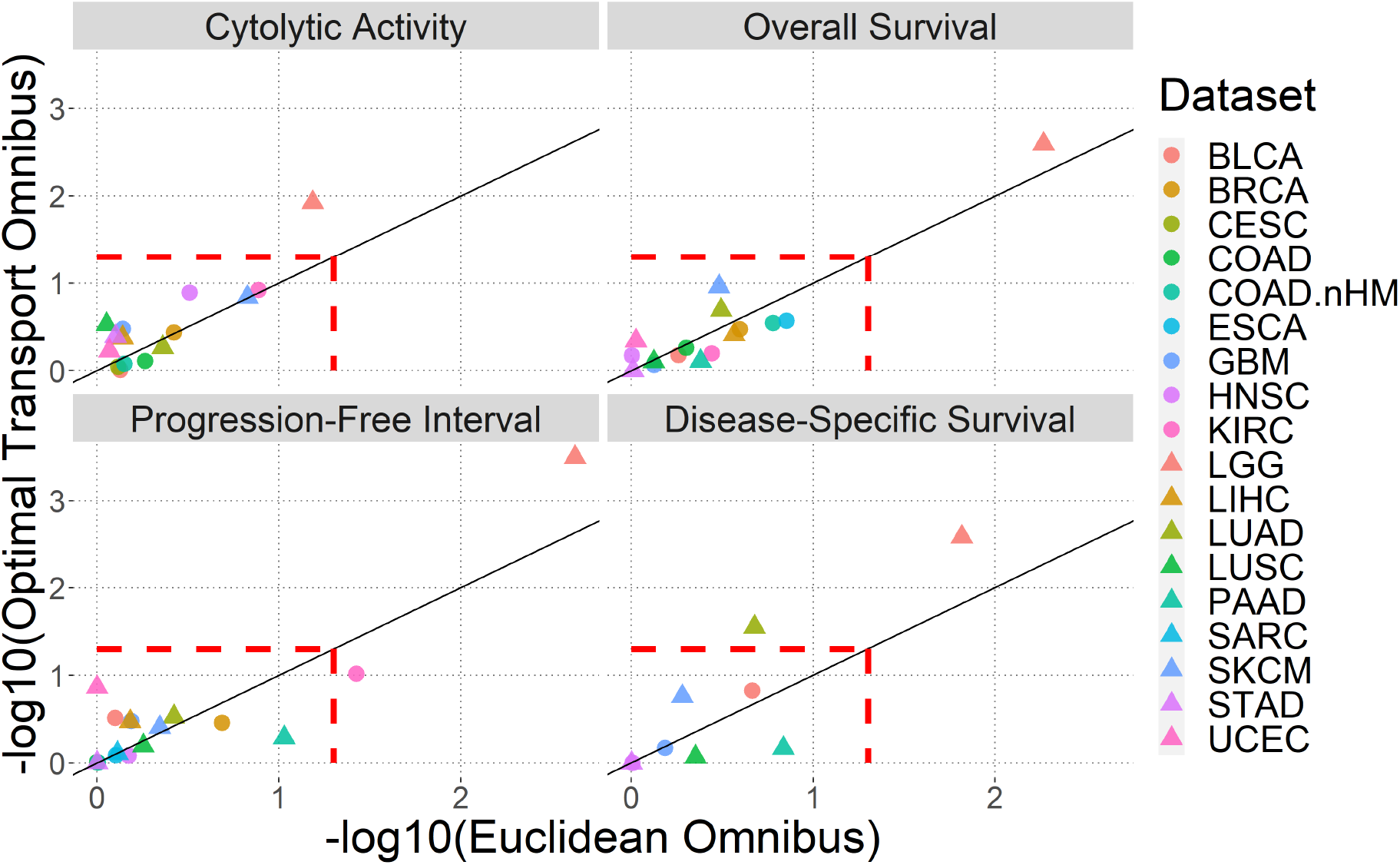
Pan-cancer kernel regression results per outcome, tumor-type and kernel approach.

We performed a backward selection to determine the null model for each cancer type and outcome. We pre-defined a set of variables to be considered as potential confounders for the null model including age at diagnosis, body mass index (BMI), BWA (alignment software version), clinical stage, gender, height, histologic grade, histologic type, binned initial pathologic diagnosis year, KIT (whole exome capture kit), log10(TMB), ploidy, purity, race, tumor stage, weight, and sequencing center. For breast cancer we also consider PAM50 subtypes obtained from Berger et al. (2018) [1]. Variables with more than 30% missing values were excluded. We performed the backward selection until no covariate had p-value greater than 0.15. log10(TMB) was not subject to variable selection and included throughout the model selection. The final null model for each cancer type and outcome is presented in Web Figure 4.

**Figure 4:**
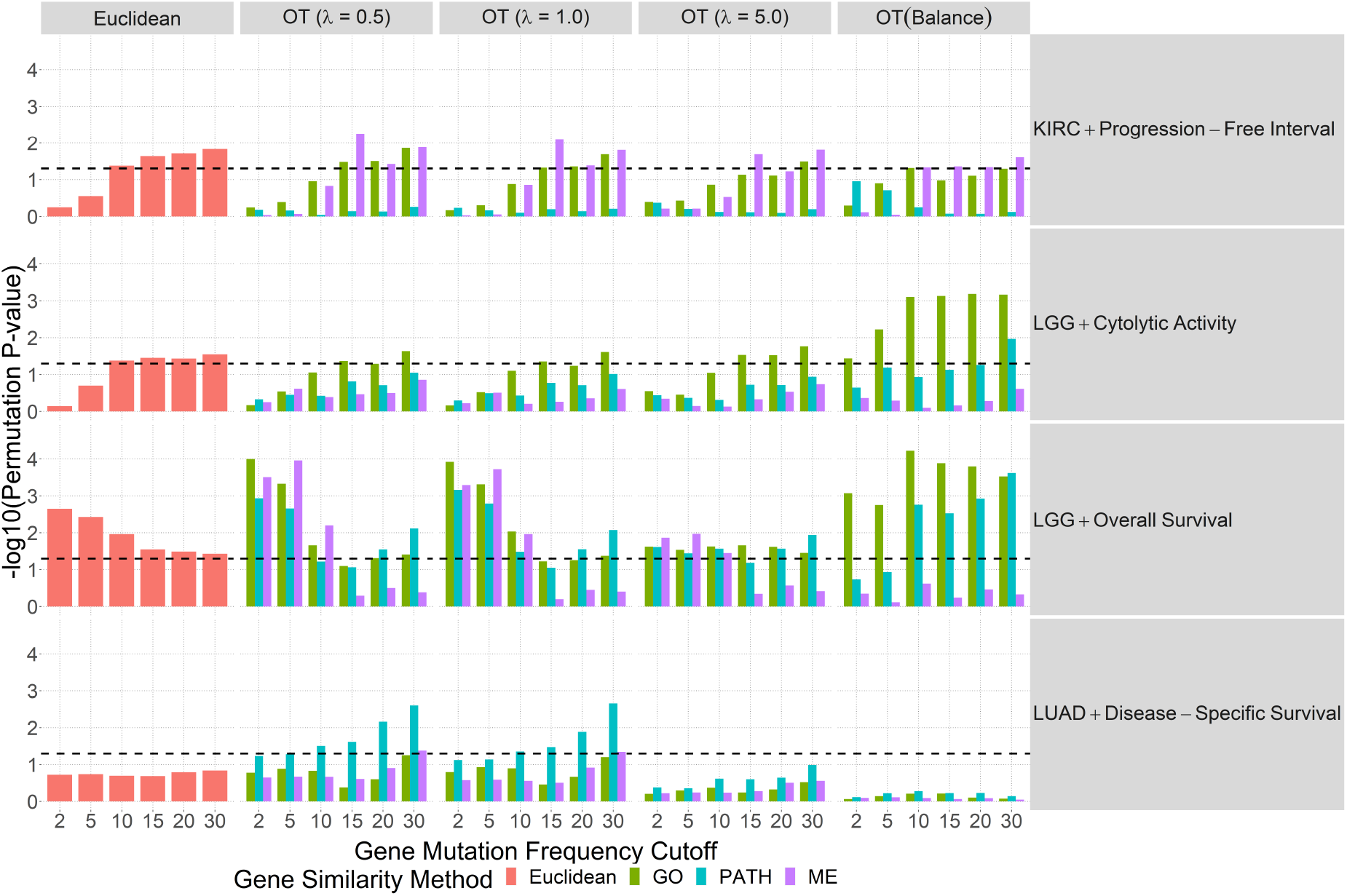
Individual kernel associations for the significant overall somatic associations with cancer types, KIRC, LGG, and LUAD.

For kernel regression, OT and Euclidean distances were transformed into positive semi-definite similarity/kernel matrices as described in Section. A total of 100,000 permutations were used to calculate the p-values. With an overview from our analyses (Figure 3), we see that most cancer type and outcomes were insignificant by both OT and Euclidean approaches with the exception of three cancer types, kidney renal clear cell carcinoma (KIRC), lower grade glioma (LGG) and lung adenocarcinoma (LUAD) for which the association between outcome and somatic mutation was significant with p-value below 0.05 with at least one outcome by either OT-based and/or Euclidean-based omnibus tests. For the complete pan-cancer results for individual Euclidean and OT kernels, refer to Web Figures 5-21.

For KIRC, only the Euclidean approach was significant for progression-free interval (PFI) when accounting for six tested kernels. Regarding individual kernels (Web Figure 12), both Euclidean and omnibus OT methods identified significant somatic mutation associations with PFI when considering the genes with mutation frequency greater than 15. For OT, significant associations were observed for both mutual exclusivity and gene ontology kernels.

For LGG, both omnibus tests by OT and Euclidean kernels were significant for all outcomes. Referring to the individual kernels (Web Figure 13), the GO-based gene similarity provided optimal significance at various mutation frequency cutoffs. The OT kernel with PATH-based gene similarity yielded smaller p-values as fewer genes were considered. Balanced OT corresponded to the individual kernel with greatest significance in all four outcomes. Based on the results, this suggests that LGG samples have similar number of disease associated pathways.

For LUAD and DSS outcome, the OT omnibus test reached the significance threshold while Euclidean omnibus did not. All individual Euclidean kernels were insignificant at various mutation frequency cutoffs while OT with PATH-based gene similarity had increasing significance for lower OT penalties and smallest individual kernel p-value at mutation frequency cutoff of 30, corresponding to about 394 genes. This suggests that the OT approach favors the variability in TMB from samples and that PATH-based similarity captures some underlying pathway. From simulation, lower penalties with a variable number of mutated pathways lead to slightly higher power. In fact, the insignificance of balanced OT across mutation frequency cutoffs, outcomes and gene similarities reinforces this possibility.

## Discussion

While we observe mutations of individual genes, these mutations often affect phenotype through perturbation of biological processes or pathways. However, it is difficult to infer the pathway-level perturbation from gene-level mutation data. We overcome this challenge by a kernel regression approach. Specifically, we ask whether patients with similar mutation data tend to have similar clinical outcomes. Our method ROKET defines the similarity of mutation data to capture the pathway level similarity, by exploiting gene annotations and a novel application of optimal transport method. Our approach aims to improve the power to detect associations over testing genes one-by-one with multiple testing correction and the biologically agnostic distance-based kernels.

The pan-cancer analysis revealed numerous insights regarding the association analysis of somatic mutations with clinical outcomes. One insight is a naive calculation of gene-gene similarity by their mutual exclusive pattern is confounded by mutation frequency and we need to account for mutation frequency by a regression approach. A second insight arose from the fact that association between somatic mutations and clinical outcomes may be only identified when considering a specific type of gene similarity definition. Lastly, the elegance of the optimal transport framework to exploit gene-gene similarity to calculate sample to sample similarity shows promise for future endeavors.

The main challenge was not knowing a priori how many genes to supply for optimal transport, how to arrive at gene similarities given existing notions and annotation, and which tuning parameters would be best. The four scenarios and multiple tuning parameters from simulation provided some insight to guide interpretations for our pan-cancer analysis. Performing omnibus testing across multiple candidate kernels does make up for the lack of prior knowledge; however, it is at the expense of power loss. Our work suggests no single kernel based on OT penalty or gene similarity is universally the best.

While it was encouraging to see the three proposed gene-gene similarities give rise to OT distances corresponding to several statistically significant kernel associations across three cancer types, we believe there are potential associations in other cancer types missed by our current work-flow. There are several possible reasons. Specifically, we only relied on protein-altering point mutations, potentially ignoring the impact of copy number alterations. We assumed at least one of the three calculated gene similarities properly characterizes gene-gene relationships. It may be that the subset of genes potentially associated with a cancer type were not well annotated. For the GO-based and PATH-based approaches, they implicitly assume gene similarities are identical for all cancer types and thus all tissue types and cell types. Another possibility is the lack of power (small effect size or insufficient sample size, number of events) to detect the association. In several cancers, undetected associations may have been captured by clinical covariates. Future studies may consider mediation analysis or associations between somatic mutations and other genomic/clinical variables.

An overlooked aspect of OT is how to handle samples with no mutated genes. This was not an issue in our analyses with only a handful of samples having no mutations, but this can potentially raise concerns in datasets with limited mutation calls (e.g. mutation data collected by targeted sequencing). Between pairs of samples without mutations, their distance should conceptually be zero. However distances between samples with and without mutation poses an issue. One approach is to introduce a single “pseudogene” and treat every tumor as having the same single pseudogene mutated once. The pairwise similarity between this pseudogene and an actual gene is zero (cost of one) while the similarity to itself is one. However, the impact of introducing this pseudogene concept would need to be explored thoroughly in simulations and real data analyses.

Our approach leaves the door open for future directions using our current software implementation. For example, OT-based distance calculations could incorporate copy number alterations and account for intra-tumor heterogeneity.

## Supporting information

Web Files

## Acknowledgements

This work is supported by the grants from the NIH (R01CA189532).

## Data Availability Statement

The data that support the findings in this paper are publicly available at the NCI’s Genomic Data Commons PanCanAtlas Publications website https://gdc.cancer.gov/about-data/publications/pancanatlas.

## Supporting Information

Web Appendices, Tables, and Figures referenced in Section are available with this paper at the Biometrics website on Wiley Online Library. The software package can be found at www.github.com/pllittle/ROKET. It is organized as an R package to run and summarize simulations. Workflow code to process and analyze real data can be found at www.github.com/pllittle/ROKET_wf.

