## Supplementary material for "ROKET: Associating Somatic Mutation with Clinical Outcomes through Kernel Regression and Optimal Transport": Web Files

#### Contents

**1 Simulation: Gene performance** **2**

**2 Null model selection, Mutation Burden, Sample Sizes** **5**

**3 Individual Kernel Associations** **7**

### 1 Simulation: Gene performance

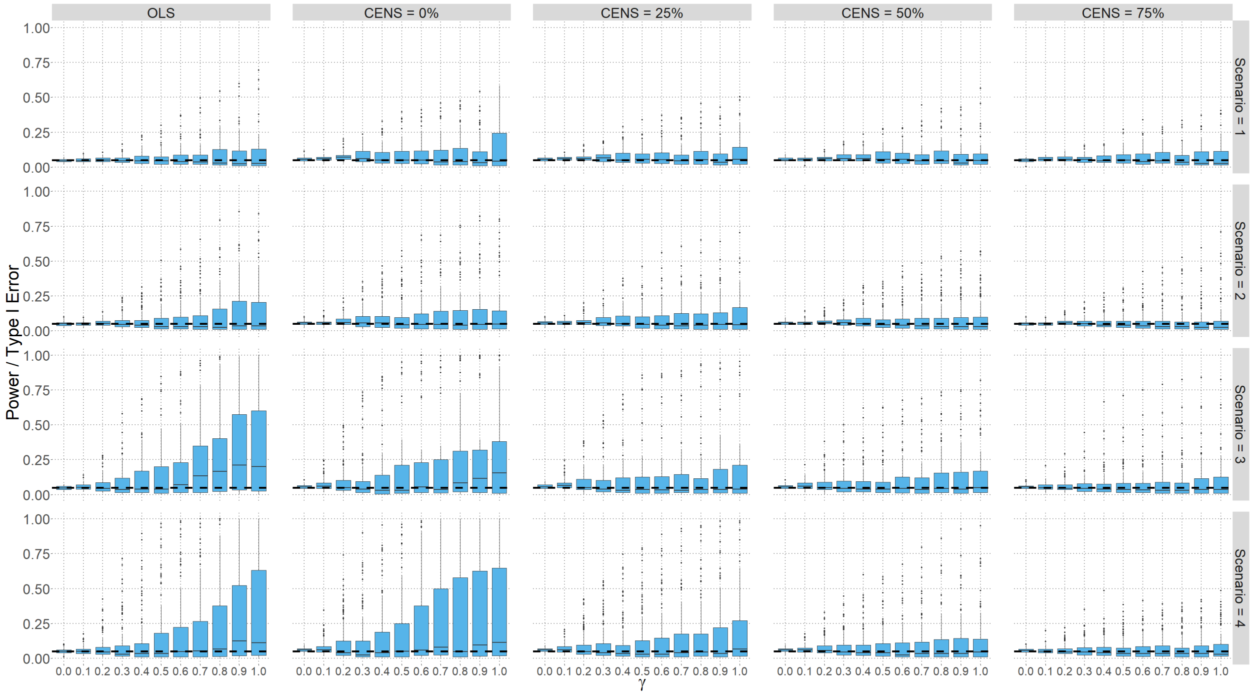


Figure 1: Summary of power and type 1 error across genes, scenarios, continuous (OLS) and survival outcomes and various censoring percentages, and effect sizes. Boxplot tick marks correspond to median proportion of rejections.

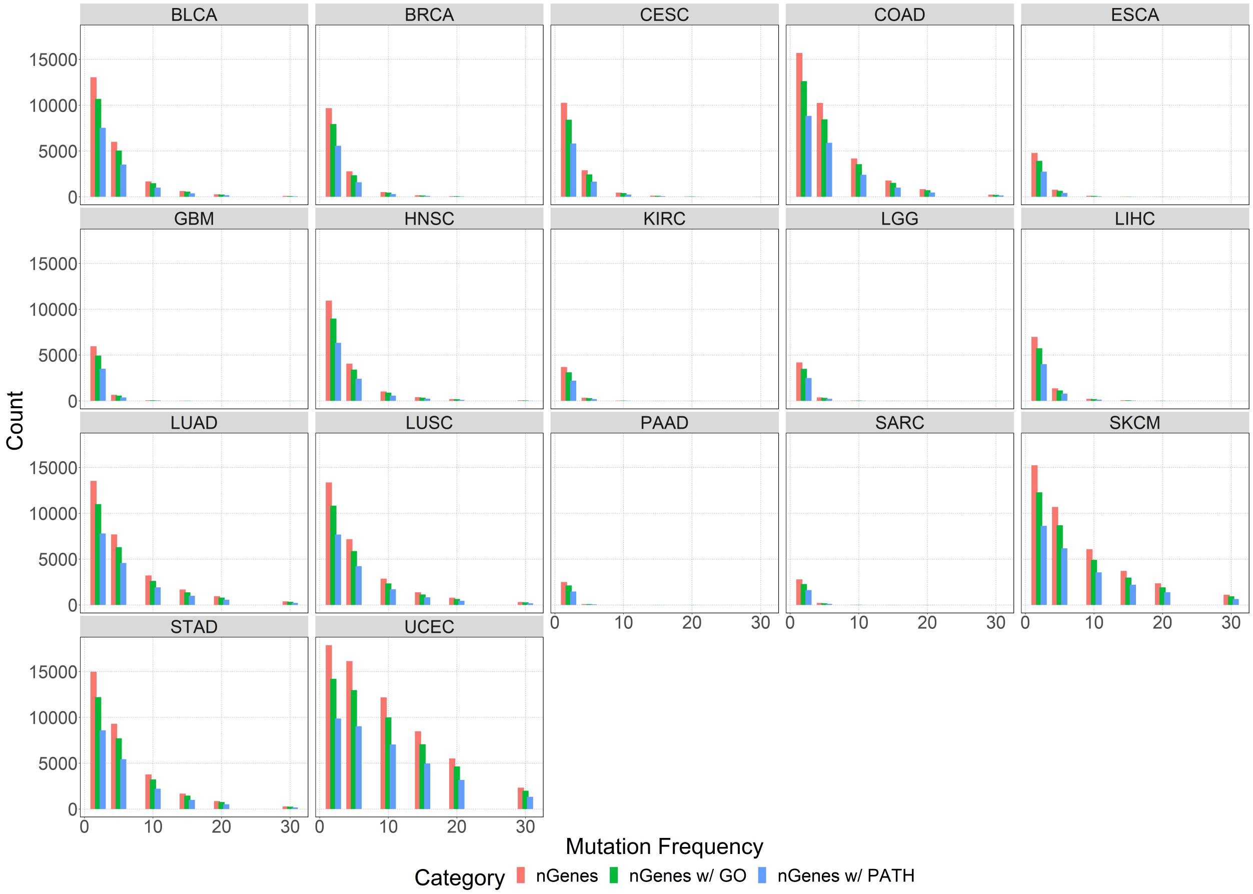


Figure 2: Number of genes with and without annotation after applying successive mutation frequency thresholds.

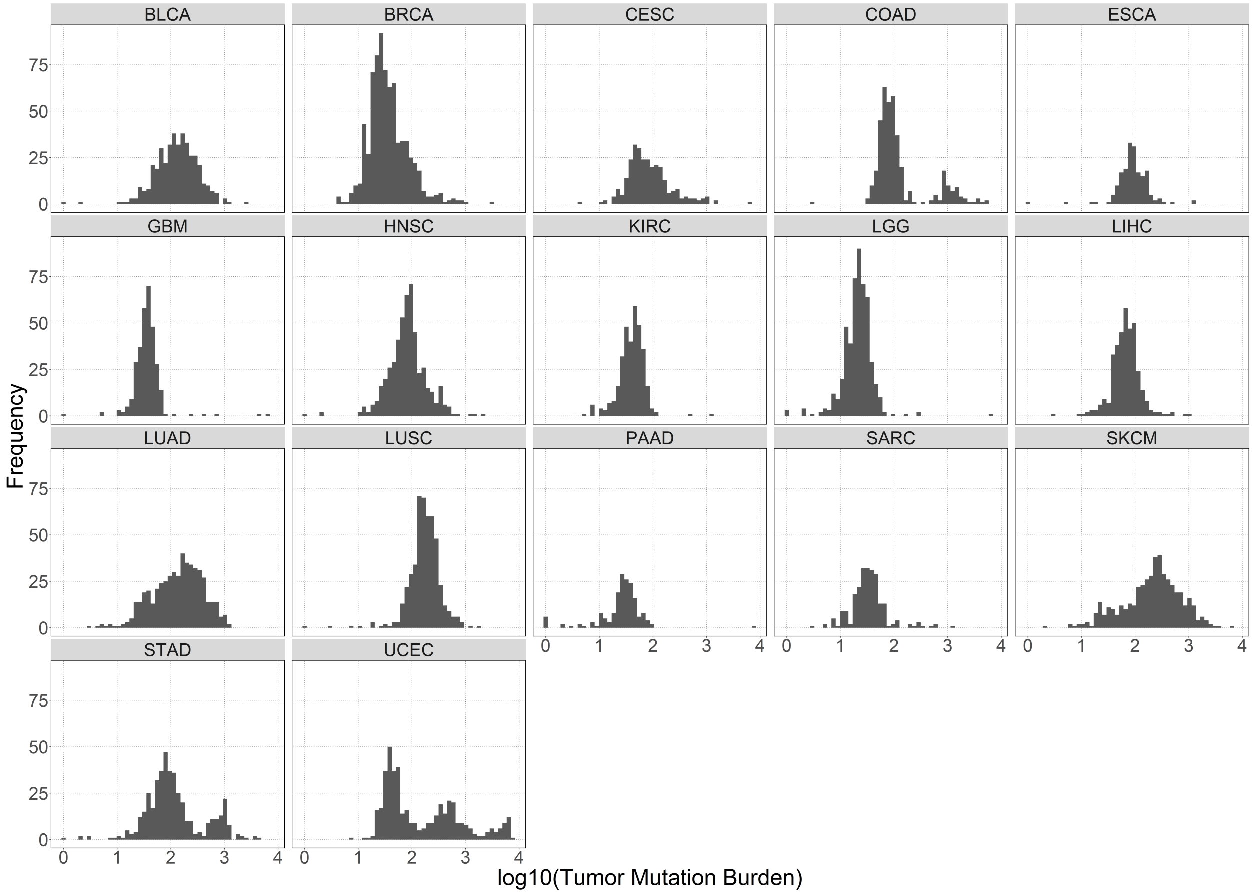


Figure 3: Distribution of log10(TMB) per tumor type.

### 2 Null model selection, Mutation Burden, Sample Sizes

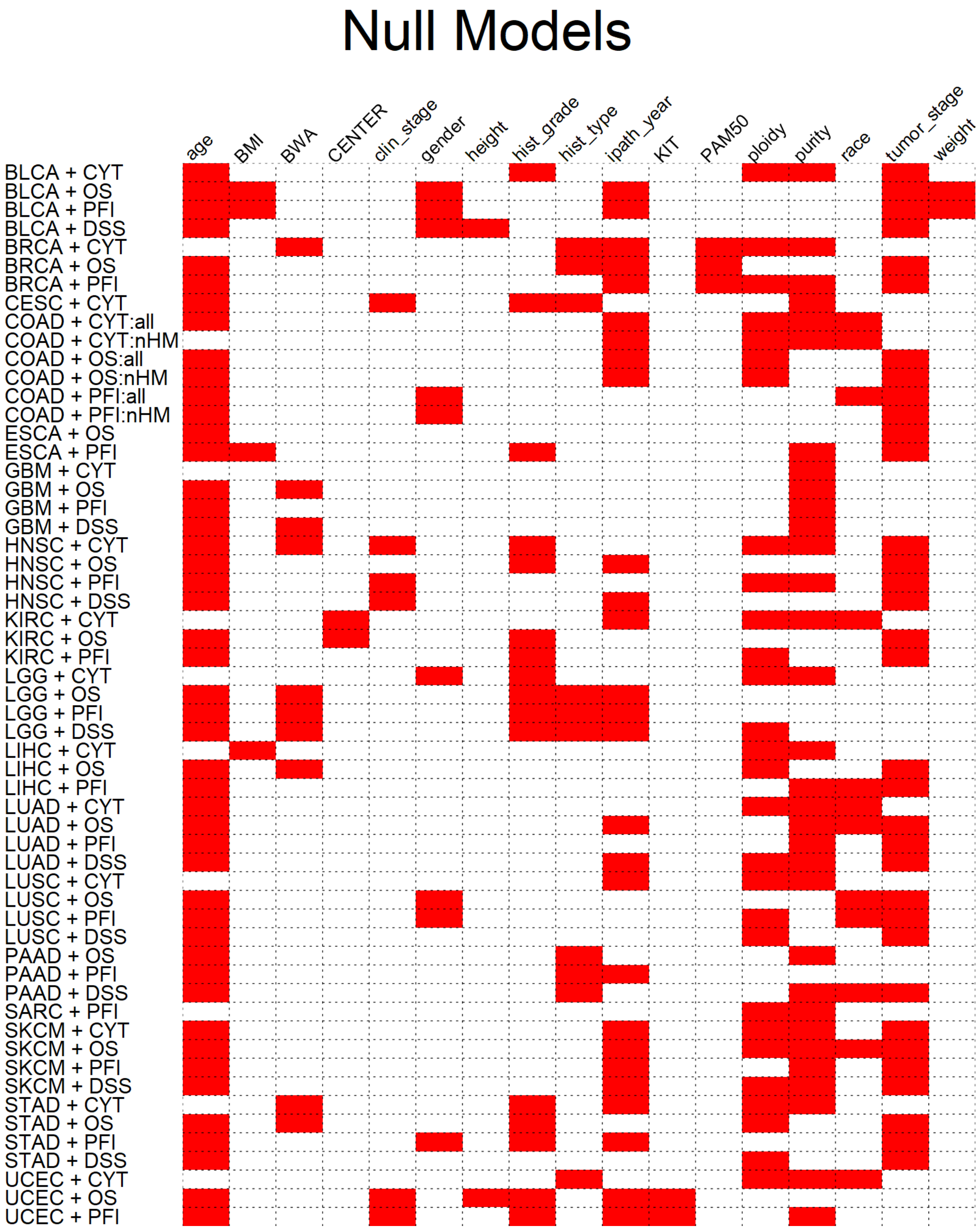


Figure 4: Null model covariates per tumor type and outcome. log10 TMB is adjusted for in all models.

Table 1: Number of subjects and number of observed events

| Name | ABBV | CYT | OS | PFI | DSS |
|---|---|---|---|---|---|
| Bladder Urothelial Carcinoma | BLCA | 240 | 341 (146) | 342 (147) | 347 (104) |
| Breast invasive carcinoma | BRCA | 576 | 639 (84) | 620 (80) | |
| Cervical squamous cell carcinoma and endocervical adenocarcinoma | CESC | 135 | | | |
| Colon and Rectum adenocarcinoma | COAD | 322 | 362 (79) | 334 (91) | |
| | COAD-nHM | 268 | 295 (65) | 299 (83) | |
| Esophageal carcinoma | ESCA | | 163 (65) | 134 (65) | |
| Glioblastoma multiforme | GBM | 112 | 217 (167) | 219 (177) | 199 (146) |
| Head and Neck squamous cell carcinoma | HNSC | 366 | 423 (184) | 402 (157) | 390 (107) |
| Kidney renal clear cell carcinoma | KIRC | 215 | 241 (65) | 234 (63) | |
| Brain Lower Grade Glioma | LGG | 317 | 438 (91) | 438 (150) | 429 (80) |
| Liver hepatocellular carcinoma | LIHC | 100 | 221 (66) | 228 (113) | |
| Lung adenocarcinoma | LUAD | 400 | 426 (150) | 481 (193) | 442 (103) |
| Lung squamous cell carcinoma | LUSC | 440 | 368 (164) | 363 (114) | 416 (81) |
| Pancreatic adenocarcinoma | PAAD | | 128 (80) | 138 (87) | 117 (59) |
| Sarcoma | SARC | | | 120 (70) | |
| Skin Cutaneous Melanoma | SKCM | 143 | 403 (190) | 410 (276) | 403 (167) |
| Stomach adenocarcinoma | STAD | 262 | 386 (148) | 397 (130) | 366 (89) |
| Uterine Corpus Endometrial Carcinoma | UCEC | 386 | 414 (70) | 426 (98) | |

### 3 Individual Kernel Associations

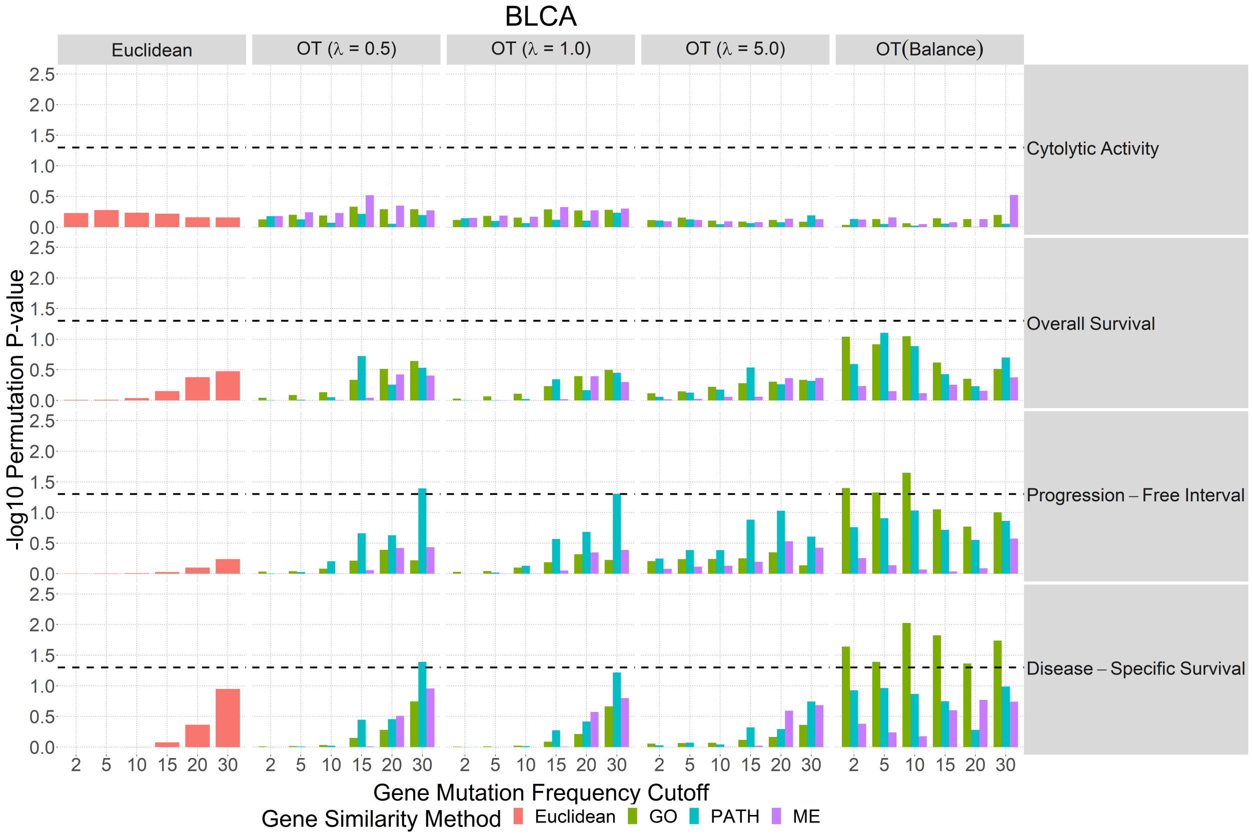


Figure 5: Bladder Urothelial Carcinoma

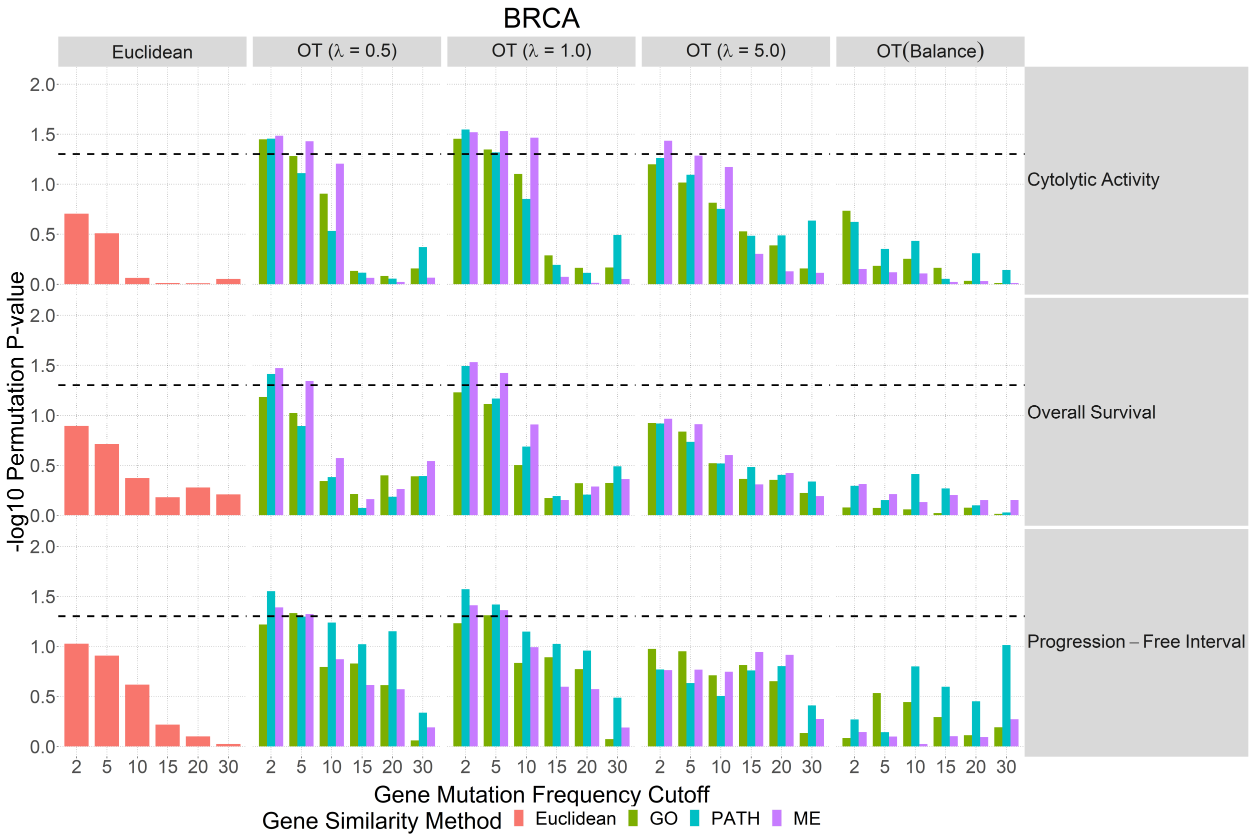


Figure 6: Breast Invasive Carcinoma

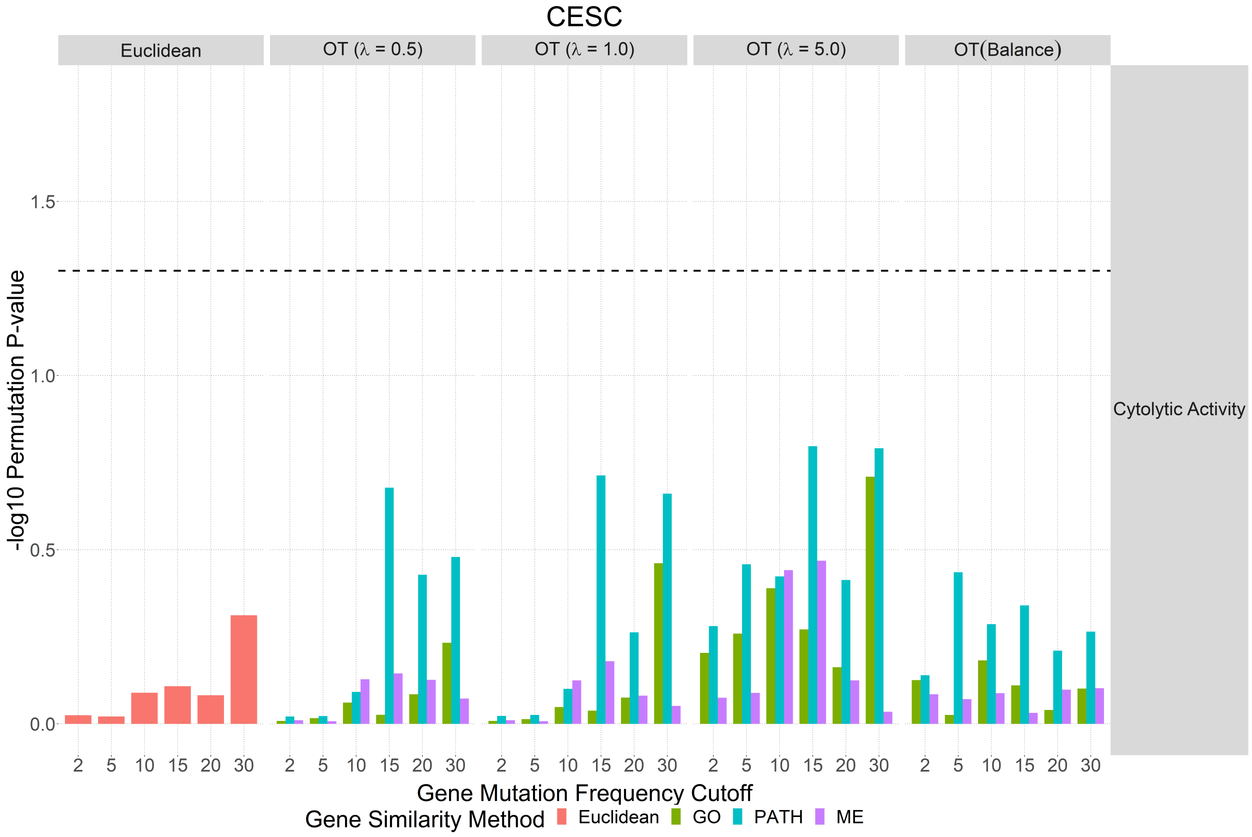


Figure 7: Cervical Squamous Cell Carcinoma and Endocervical Adenocarcinoma

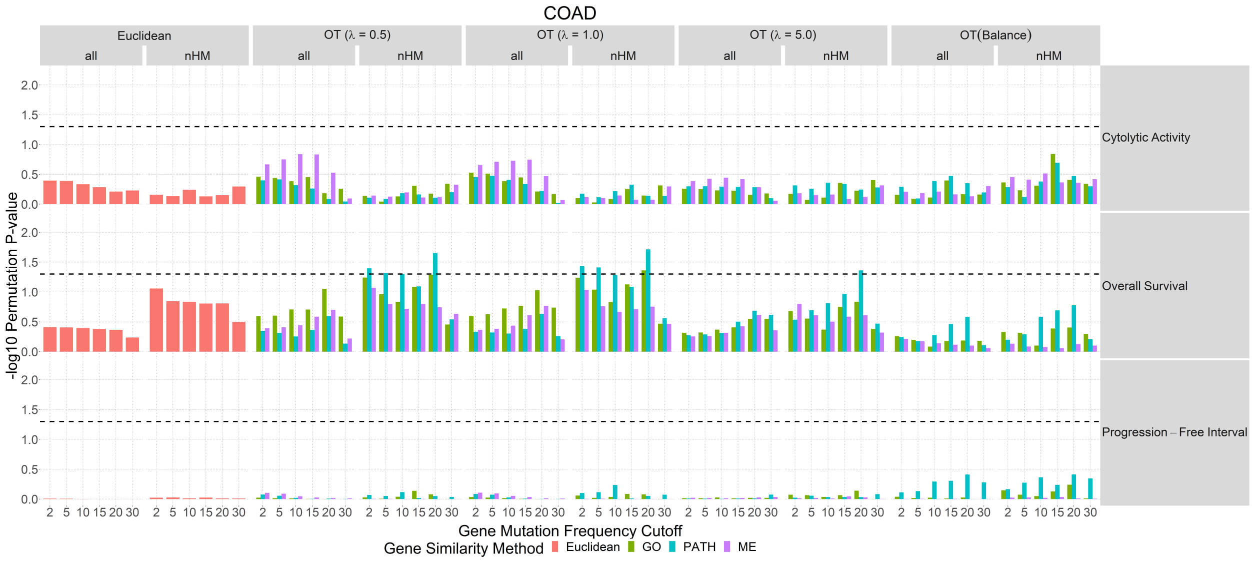


Figure 8: Colon and Rectum Adenocarcinoma

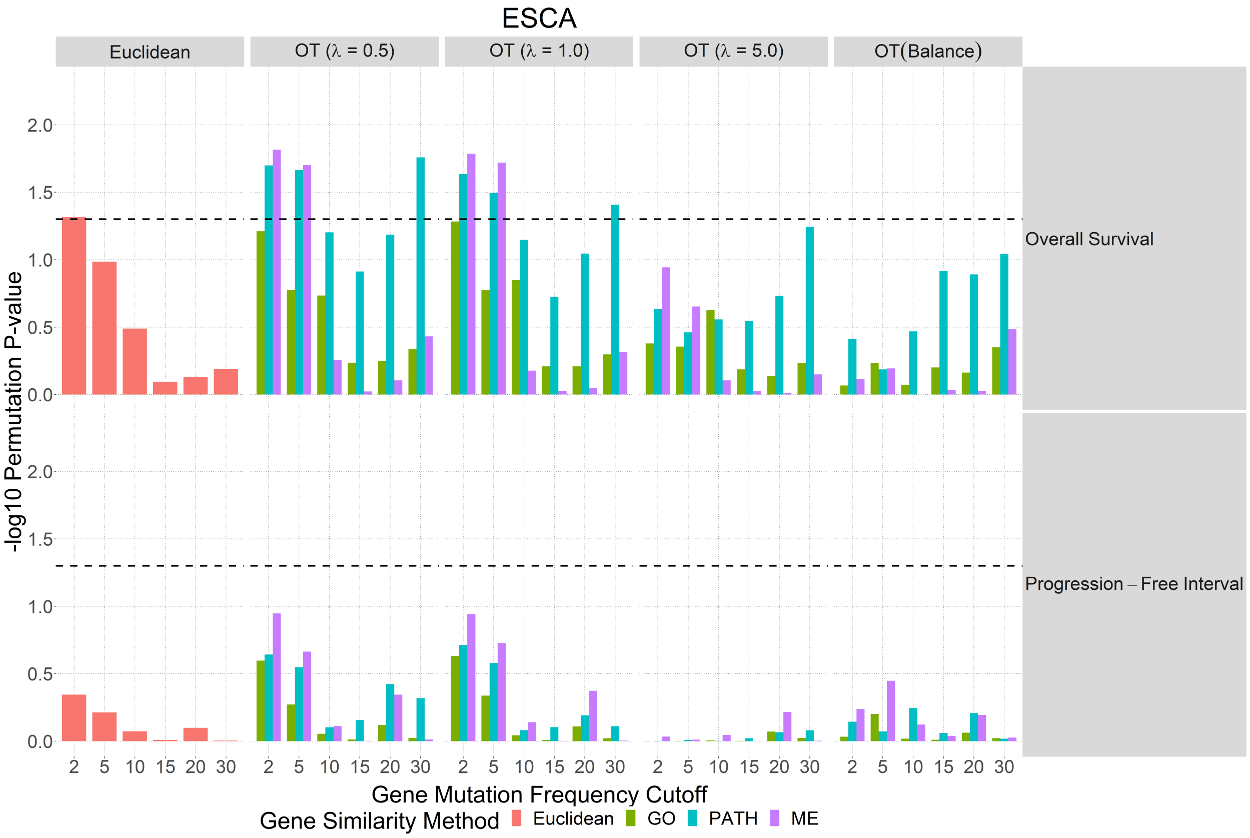


Figure 9: Esophageal Carcinoma

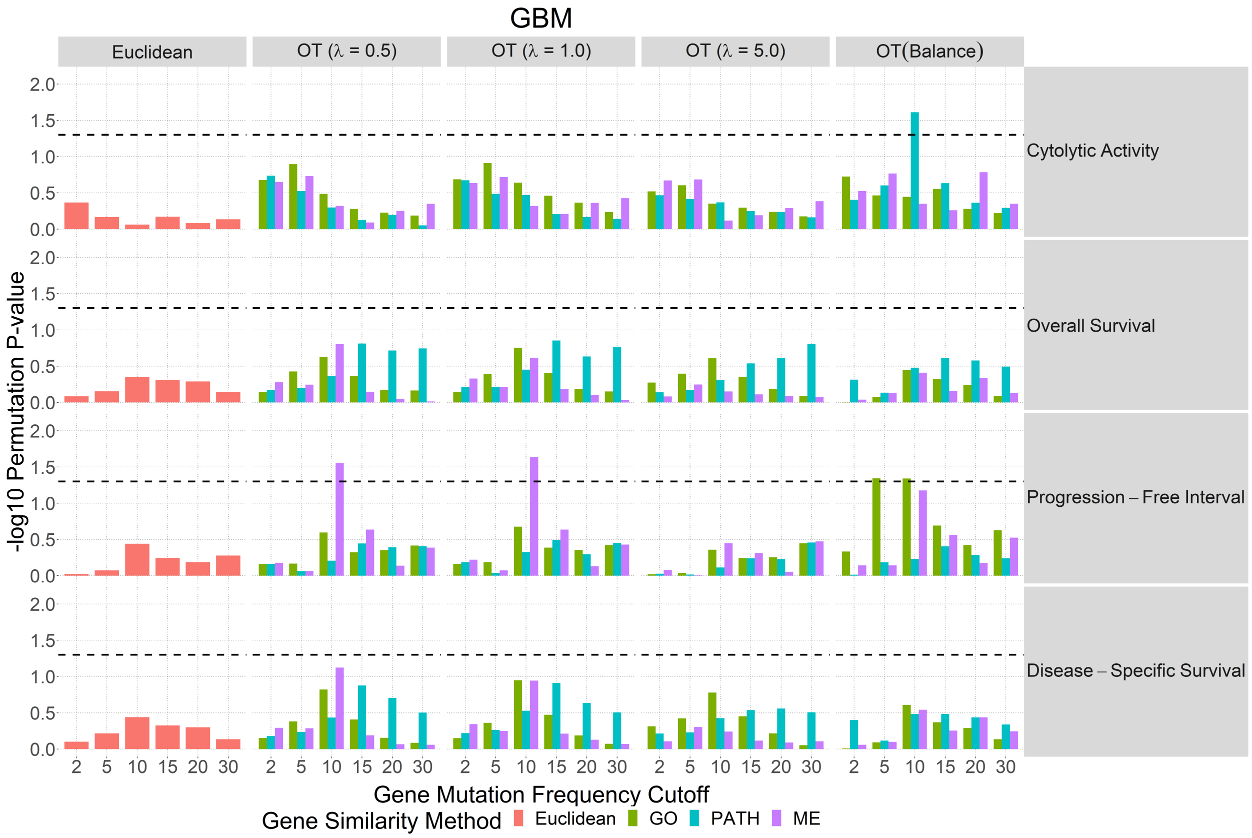


Figure 10: Glioblastoma Multiforme

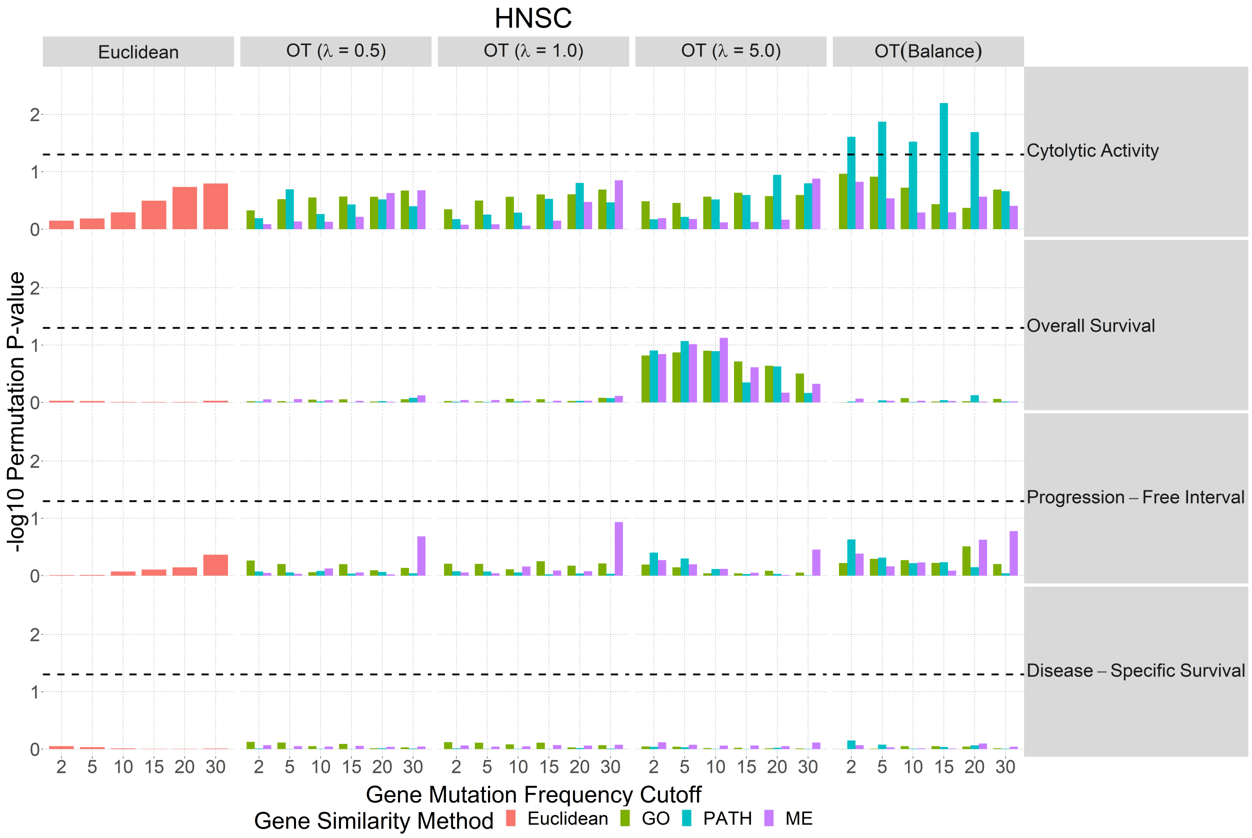


Figure 11: Head and Neck Squamous Cell Carcinoma

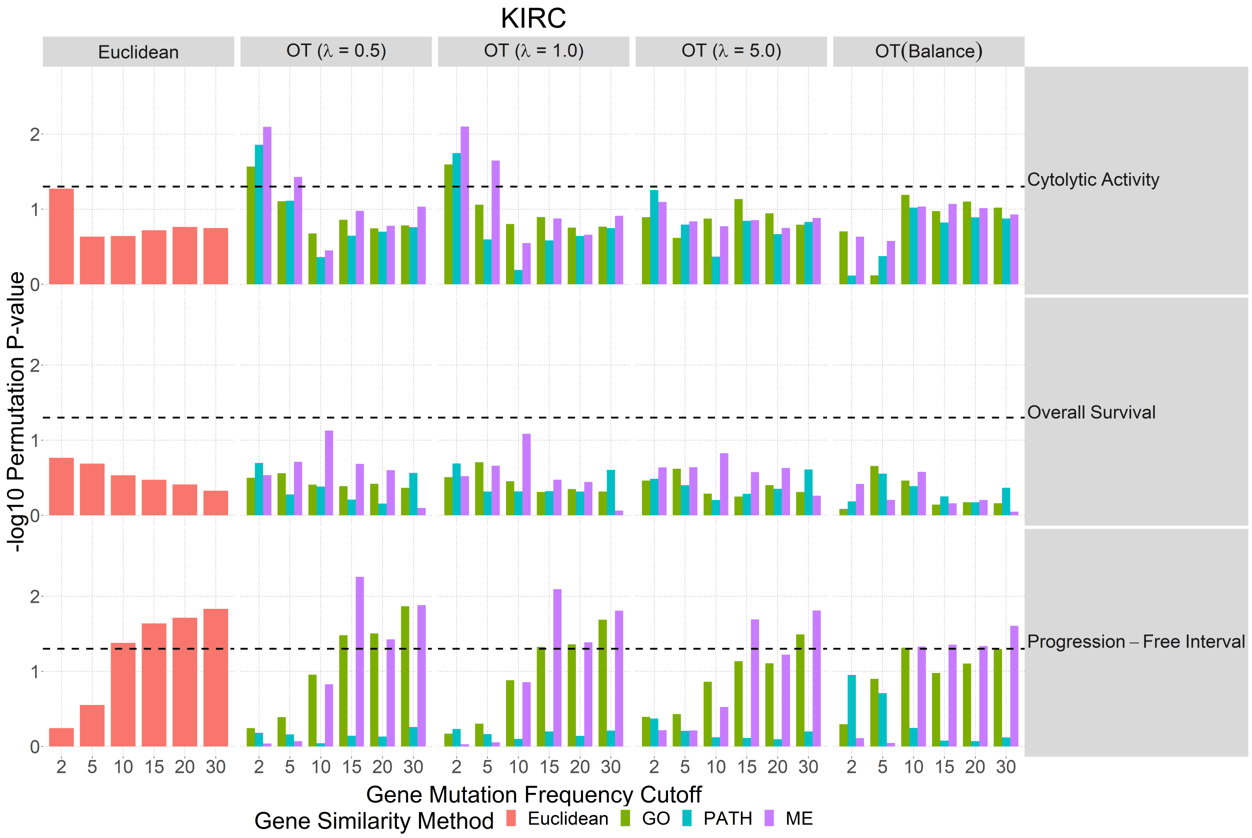


Figure 12: Kidney Renal Clear Cell Carcinoma

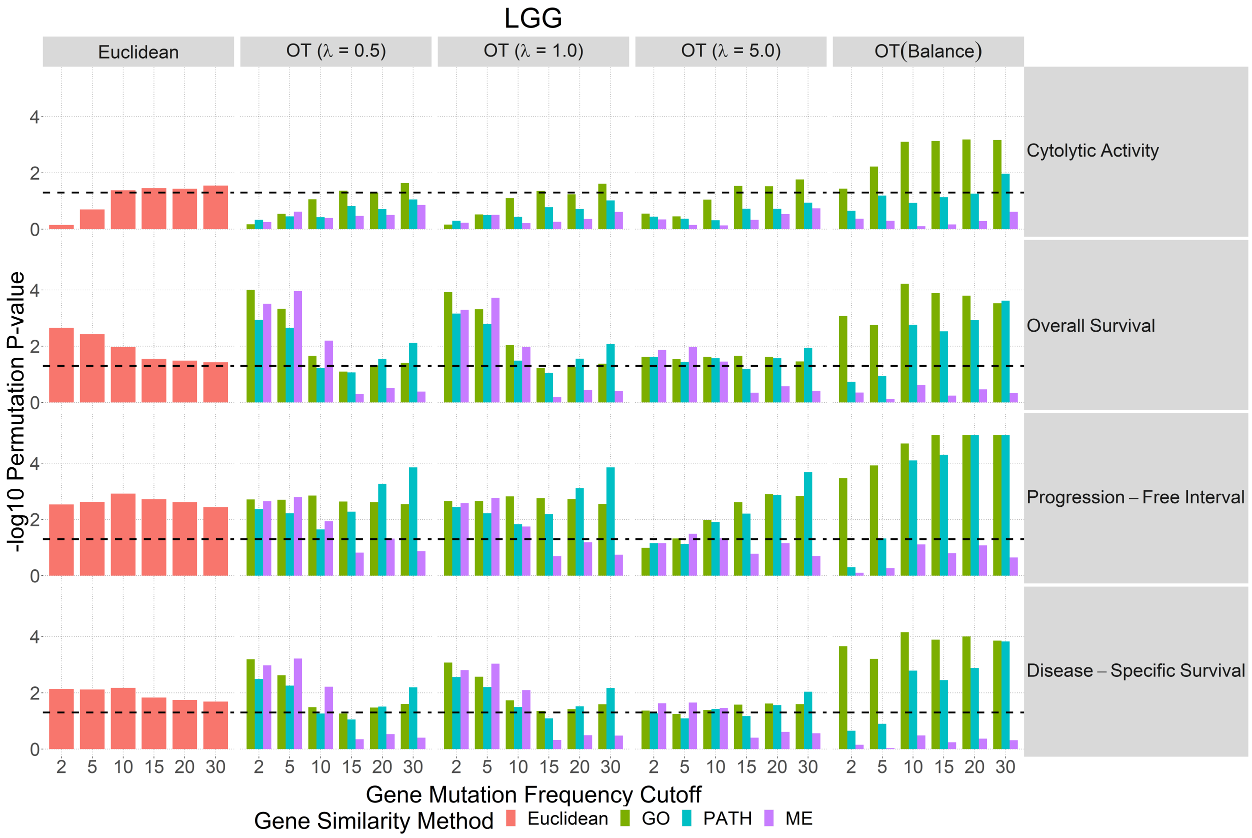


Figure 13: Brain Lower Grade Glioma

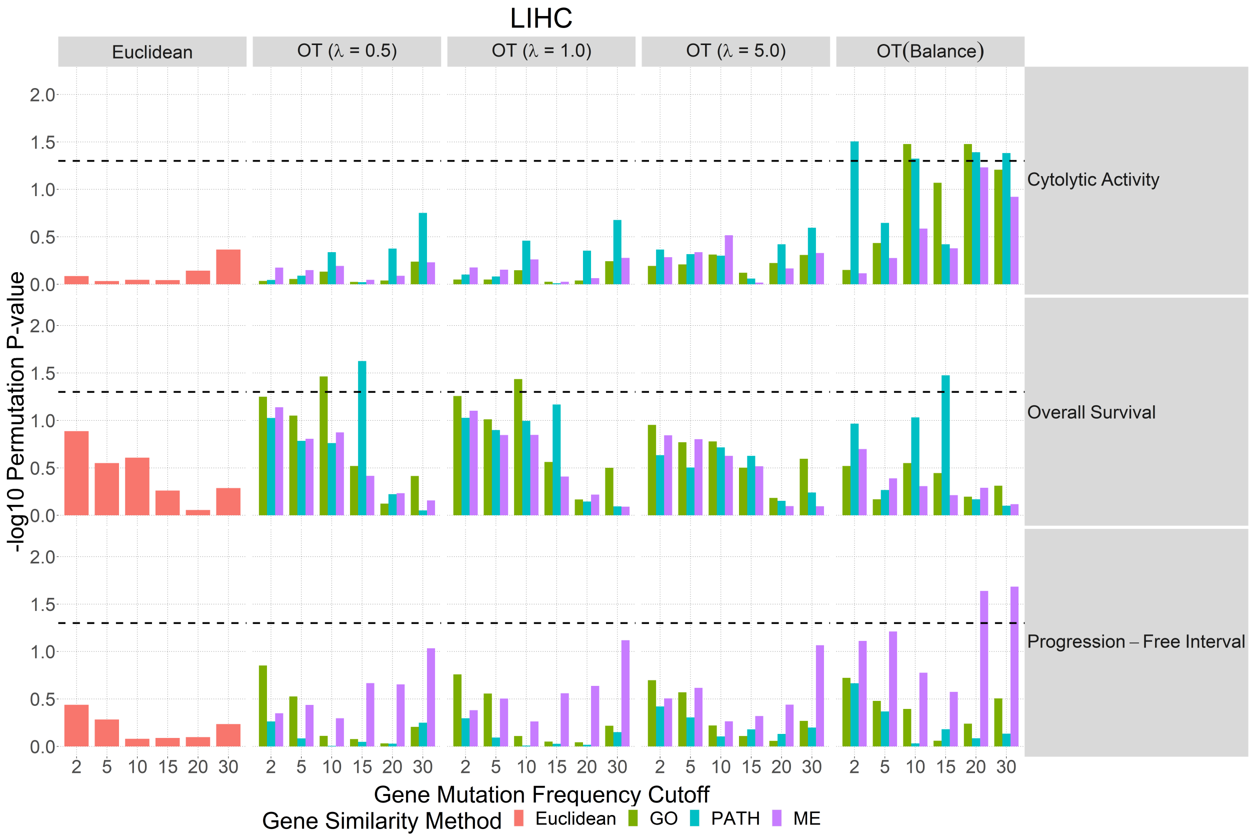


Figure 14: Liver Hepatocellular Carcinoma

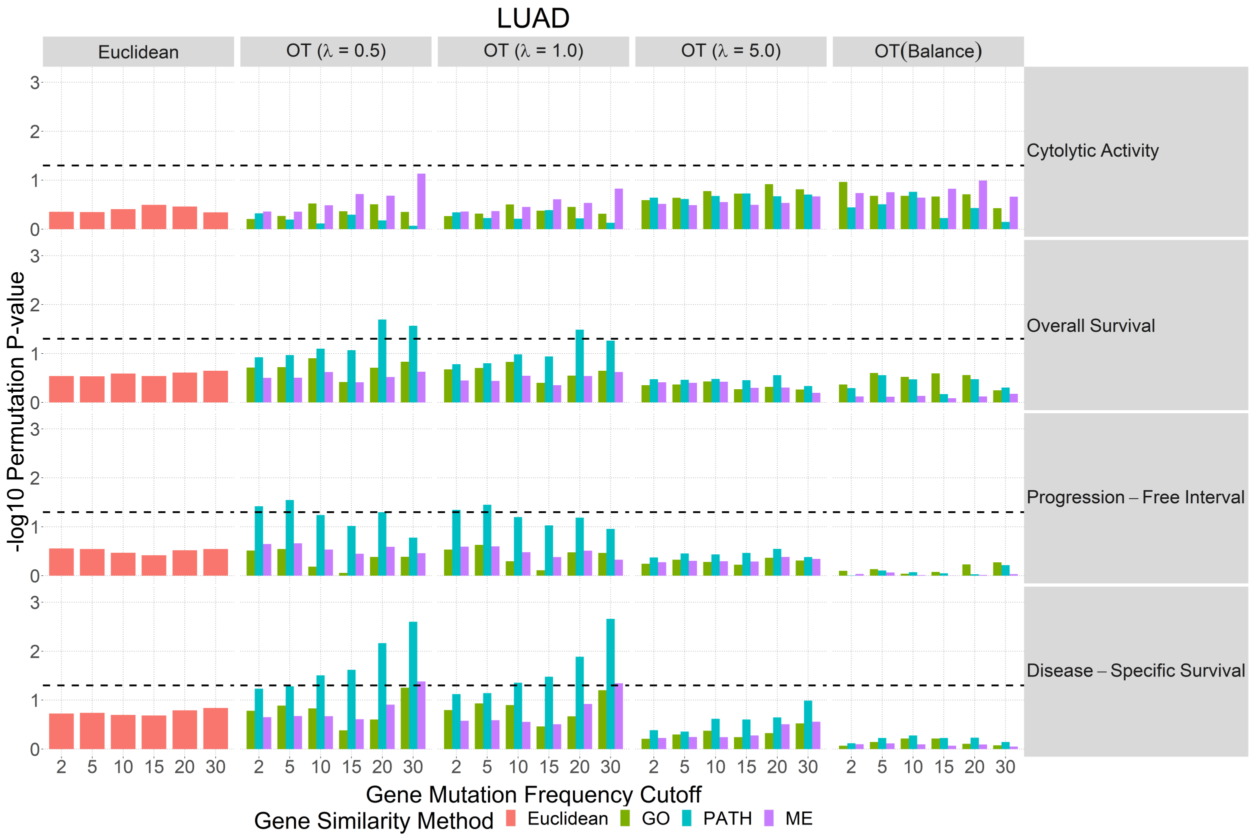


Figure 15: Lung Adenocarcinoma

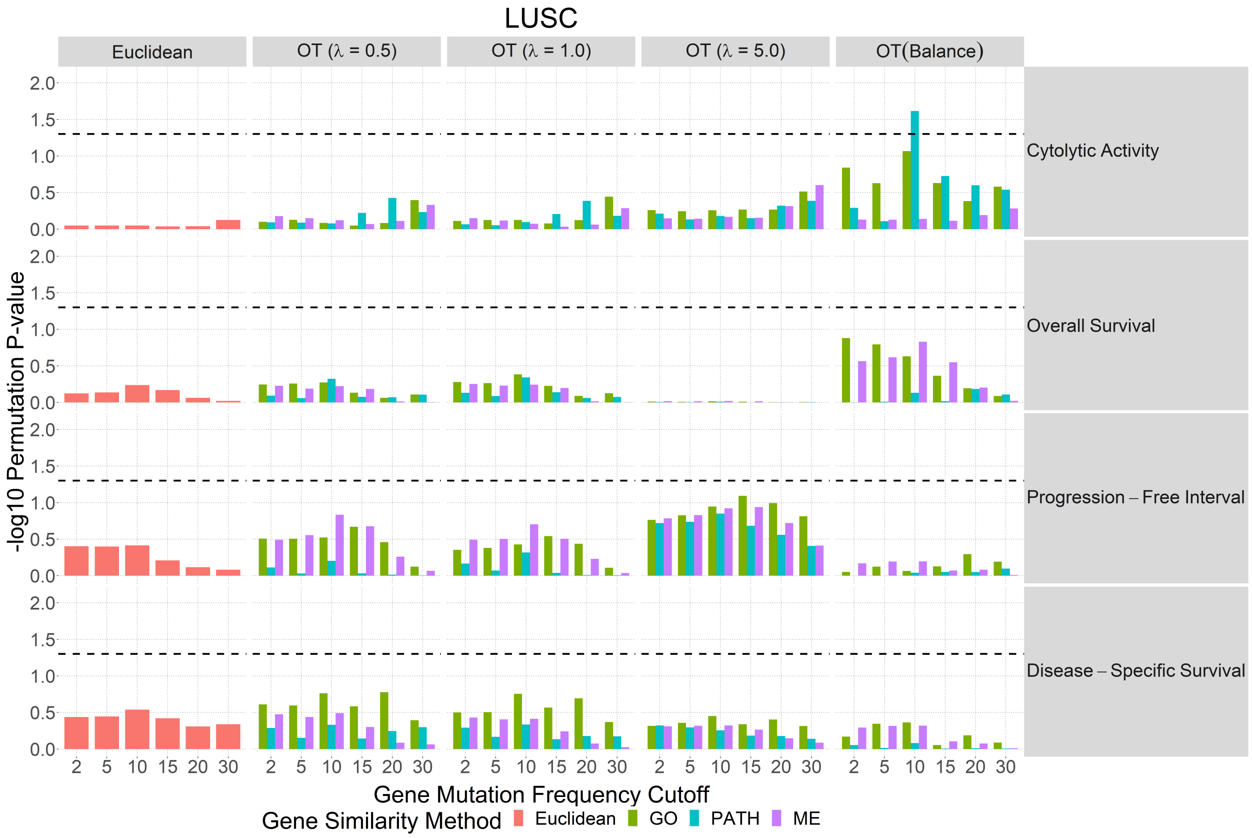


Figure 16: Lung Squamous Cell Carcinoma

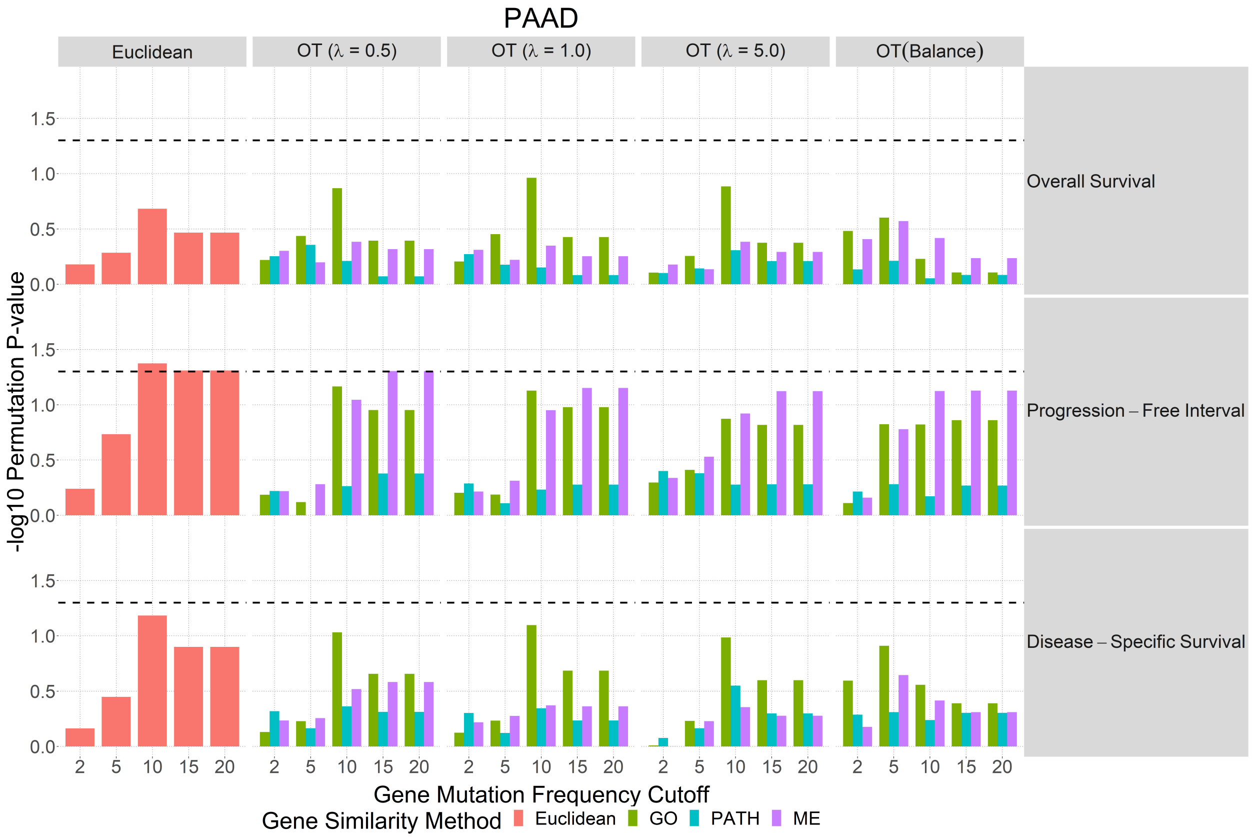


Figure 17: Pancreatic Adenocarcinoma

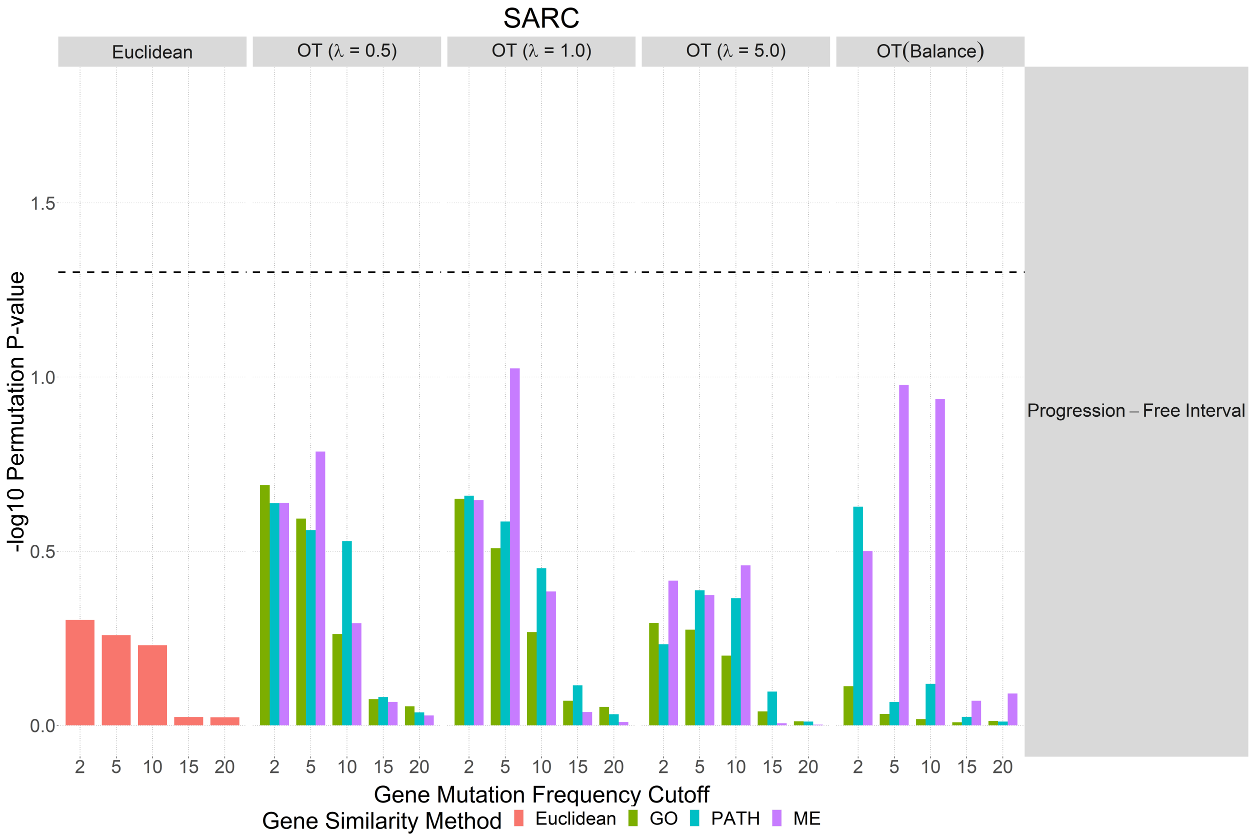


Figure 18: Sarcoma

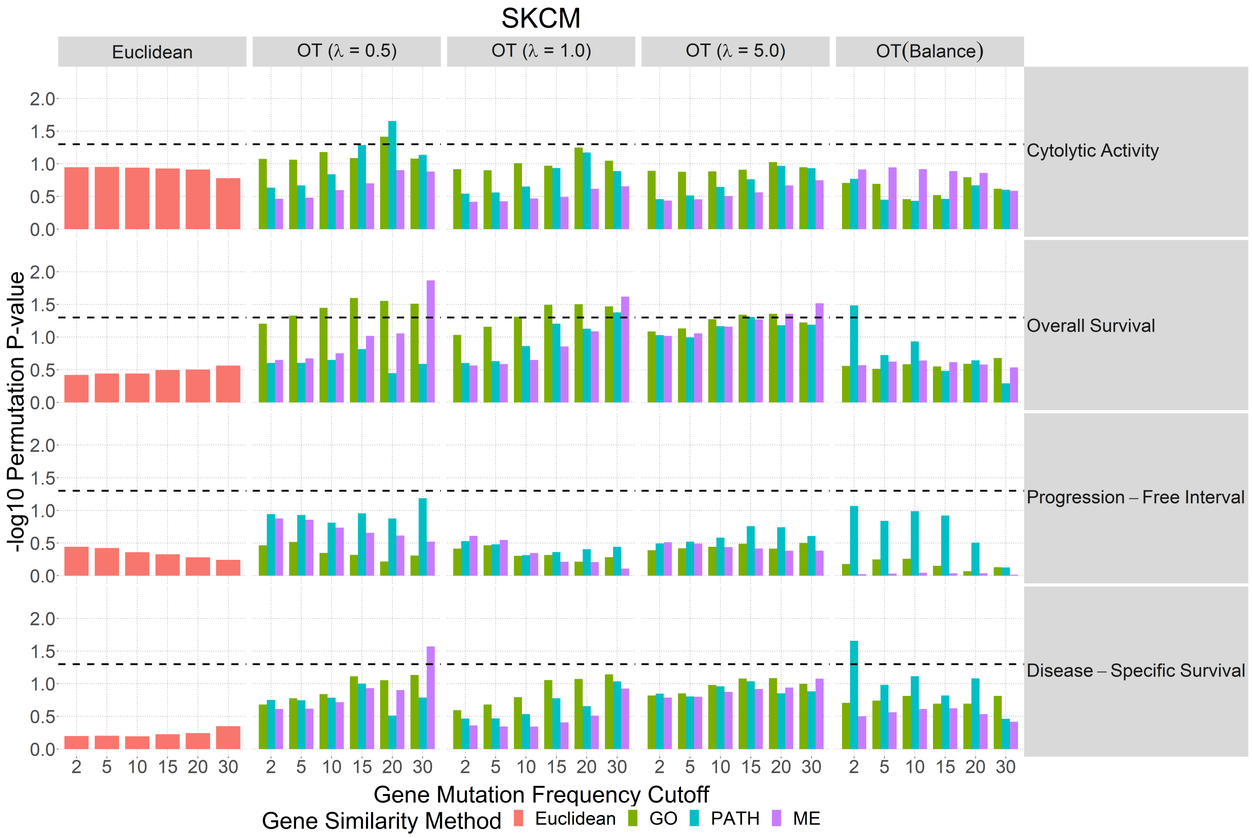


Figure 19: Skin Cutaneous Melanoma

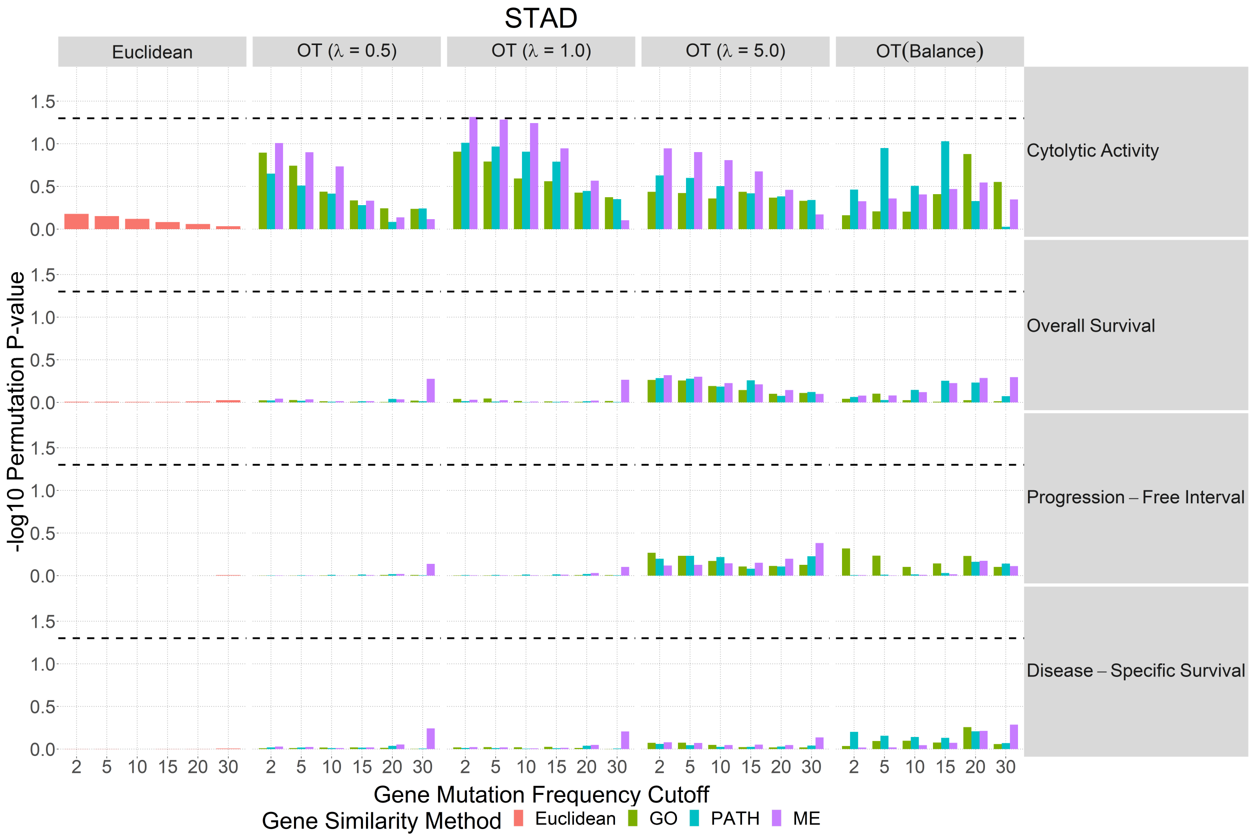


Figure 20: Stomach Adenocarcinoma

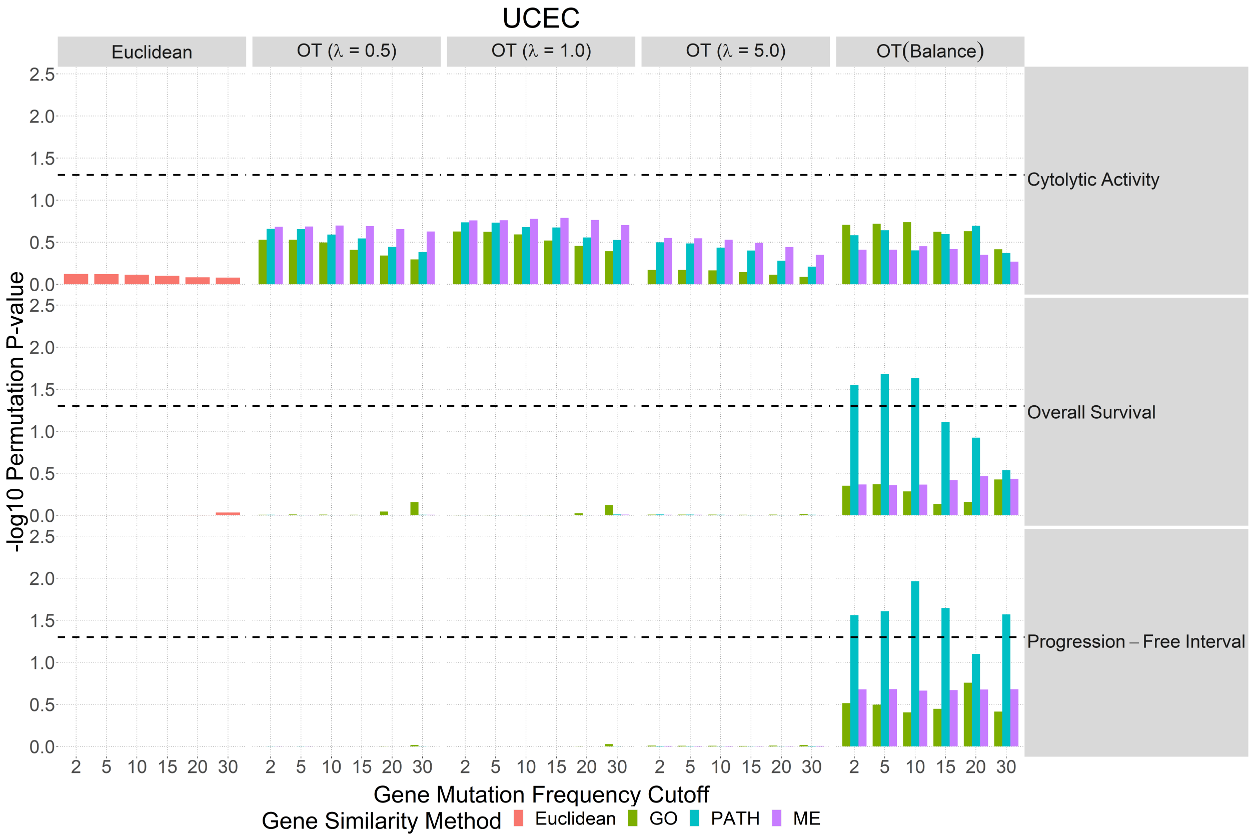


Figure 21: Uterine Corpus Endometrial Carcinoma
